# Evolutionary Stratification of Codon Usage Bias In Plants Arises from GC3 Composition and Translational Optimization

**DOI:** 10.64898/2026.06.26.734692

**Authors:** Tapan Kumar Mohanta, Sujogya Panda, Huseyin Tombuloglu, Prashant Kaushik, Jitesh Kumar

## Abstract

Codon usage bias is a universal property of genomes, yet its evolutionary bias and functional relevance across the plant lineage remained poorly defined. Here we analyzed the plant codon usage bias at the genome scale across 262 plant species, encompassing more than 4.6 billion codons from over 11 million coding sequences. Codon usage was strongly non-random and exhibited a clear phylogenetic structure, with higher plants favoring A/T-ending codons and early-diverging lineages enriched in G/C-ending codons. Integrated analysis of synonymous codon usage, codon adaptation index, and codon-anticodon pairing indicates that translational selection is mediated by tRNA availability, which contributes to the sustainability of these patterns. Machine-learning (ML) approaches identified a small subset of codons having outsized influence on genome-wide codon usage landscapes. Furthermore, we also revealed robust inverse relationships between the effective number of codons and GC content at synonymous third positions. Neutrality analysis revealed that approximately 61% of variation is driven by mutational pressure, tempered by selective constraints. Phylogenetic reconstruction showed a progressive relaxation of codon bias from algae to angiosperms, while maintaining conserved translational genetic cost of ∼ 30 ATP per codon across the lineages. Together, these results showed codon usage is an evolutionarily conserved yet lineage-specific feature that links mutation, selection, and translational optimization across the plant kingdom.

## Introduction

The genetic code, called the molecular language of life, translates the respective cognate amino acids to make a functional protein ^1–3^. The code secretly hides the biochemical and genetic language in a sophisticated way in the gene called a ‘codon’ ^4–6^. In a philosophical context, a codon can be called a symbol of intent and cosmic dialogue between chemistry and consciousness of genetic information. There are 61 sense codons, and they accommodate 20 amino acids, showing their fascinating linguistic understanding through degeneracy ^7,8^. The presence of a systematic order of codons in a gene to translate its protein is intriguing in mind, whether the codons are governed by chance or by purposeful meaning with order ^3,9,10^. Codon usage is the process in which genes are expressed in a particular organism in a specific order.

It is important to understand whether the codons are linked to each other randomly or with some meaningful context ^10–12^. The codon usage bias states a non-random use of synonymous codons that code for the same amino acid ^9^. Theoretically, each codon should get equal opportunity to be encoded in a gene. But evidence shows certain codons are preferred more frequently within a gene or a genome ^13,14^. The preferences also vary among different genes of the same organism ^14,15^. This asymmetry arises due to multiple factors like abundance and availability of tRNA ^16^, gene expression level ^17^, translational accuracy, speed, mutation pressure, selection pressure, and drift ^14,18–20^. Shannon’s information theory defines redundancy as a measure of predictability, and it can be called a biological compassion ^21–23^. The bias mirrors the debate between determinism and free will. The “Leibniz’s principle of sufficient reason” explains that nothing exists without a cause ^24^. Therefore, the codon bias is not random but a consequence of adaptive requirements. Further, a question arises: why are certain codons preferred over others when both can encode the same amino acid? Do codon optimization and performance play a role in such selective codon usage? Or does the genome choose a suitable codon in the concept of molecular economics to save energy? The study regarding the extension of the genetic code with a quadruplet anticodon and the report on Hachimoji DNA ^25^ with extended base pairs shows codon usage bias is not just a natural phenomenon but rather a sophisticated design parameter. Across the tree of life, codon usage shows extensive evolutionary conservation and divergence. Organisms with similar functionality sometimes share codon preferences explaining their functional convergence. Therefore, we conducted a genome-wide codon usage bias analysis of 262 plant species and examined the results in detail.

## Material and Methods

### Sequence downloads and codon usage

The CDS sequences of the plant species were downloaded from the National Center for Biotechnology Information (NCBI) and the Phytozome databases ^26,27^. Subsequently, the codon sequences were subjected to codon usage analysis using a Python script. The number of codons, frequency, and relative synonymous codon usage were recorded for further analysis. The plant species with codon information were grouped according to their lineages for further analysis. All the files associated with the codon usage were kept as comma-separated (.csv) files.

### Codon annotation and relative synonymous codon usage

Codon usage data from multiple plant species were converted to proportions before their analysis to ensure numerical consistency across downstream computations. The codons were mapped to their corresponding amino acids using the standard genetic code. The stop codons were excluded from the study. The relative synonymous codon usage (RSCU) was calculated for the codons using the standard genetic code, where RSCU was the observed count of codons divided by the mean frequency of all synonymous codons encoding the same amino acid. An RSCU > 1 indicates preferential codon usage, while a value < 1 indicates avoidance^9^. The number of preferred synonymous codons with RSCU > 1 was quantified for each species, and the resulting values were combined with the lineage metadata to generate the lineage-wise result of the codon preference pattern. The lineage level mean relative synonymous codon usage was calculated by averaging RSCU values across all species within each lineage.

### Codon Adaptation Index (CAI)

For the codon adaptation index ^28^, a reference codon usage table from the top 100 highly expressed genes of *Arabidopsis thaliana* was used to represent the optimal translational preference, combined with the codon usage frequency of multiple species. The stop codons were excluded, and only the sense codons shared between the files were retained for the study. The codons were assigned as per the standard genetic code, and the synonymous codons were grouped to allow normalization within each amino acid family. The relative adaptiveness weights were calculated from the reference dataset by dividing the frequency of each codon by the maximum frequency observed among synonymous codons encoding the same amino acid. This assigns a weight of 1 to the most preferred codon for each amino acid. The CAI values were later computed for each species as the weight of the geometric mean of codon adaptiveness values by using genome-wide frequencies as weights. This excluded the codons with zero frequency or undefined weights. Species that lacked informative codon contributions were assigned a missing CAI value. The differences in CAI between the lineages or species were evaluated using one-way ANOVA and post-hoc pairwise comparison with Tukey’s honestly significant differences (HSD) test to identify significant differences in translational adaptation (p < 0.05). The resulting CAI values were summarized across the species, and the results were visualized in histograms. All analyses were performed using the Python-based program.

### Codon-Anticodon Mapping

To map the codon-anticodon frequency, first genome-wide anticodon data of the plant species were used, which were generated previously ^29^. Codon usage and anticodon usage data were integrated for subsequent analysis. All frequency values were converted from percentages to proportions prior to the analysis. The codons and anticodons were mapped according to the standard A-T and G-C base pairing, and a codon-specific tRNA availability table was constructed as the proportionate abundance of the cognate anticodon. Codon that failed to match the corresponding anticodon were assigned zero. The generated matrix was cleaned where the codons lacked a detectable tRNA signal across all samples. To visualize the global patterns of codon-anticodon correspondence, a hierarchical clustered heatmap was generated using a Euclidean dissimilarity matrix and average linkage for cluster aggregation. Subsequently, cophenetic correlation coefficients were calculated to quantify the robustness of the analysis and confirm the preservation of pairwise distances. To understand the overall codon usage bias, raw species were standardized by z-score normalization. The color intensity depicts the relative availability of tRNA. The cluster patterns were evaluated statistically by comparing within-versus between-cluster variance using one-way ANOVA with tRNA-codon association at *p* < 0.05. All the statistical analyses and visualizations were conducted in a Python-based platform that used the core Python language, CodonW, tRNAscan-SE, and the Pandas library.

### Molecular Economy of Codons

The molecular economy of plant codons was studied by integrating the codon usage data with the nucleotide biosynthetic cost as ATP equivalents ^30,31^. Prior to the analysis, the percentage frequency of codon usage was converted to proportions and the biosynthetic cost of each codon was calculated as the sum of the ATP equivalents of the nucleotides (A:12, T:9, G:12, and C:8) ^31^. To determine the molecular economy of the plant species, we summed up the cost of each codon, weighted by its frequency of the codons across the genome. This gave us an idea of how much energy each codon cost in translation. The molecular economy distribution across the species was visualized in a histogram with kernel density estimation to assess skewness and multimodality. Statistical differences in molecular economy between the predefined groups or lineages were evaluated using one-way ANOVA with a post-hoc Tukey’s honest significance difference (HSD) test to identify significant differences in translational energy efficiency. All the studies were performed using Python-based pipelines.

### Effective Number of Codons (ENC)-GC3 Calculation

Genome-wide codon usage bias across multiple species was calculated using the effective number of codons (ENC), GC content at the third position (GC3), and relative synonymous codon usage (RSCU) from the codon frequency table. The codons were assigned to their respective amino acids and grouped into the degeneracy class (2, 3, 4, and 6 folds) to enable the F_k_ statistics for ENC ^32^. The F_k_ statistics measures how unequally the synonymous codons are used 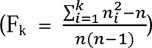^32^. The F_k_ statistics of each species were calculated as the weighted sum of F_k_ values. This provides a quantitative measure of genome-wide codon usage bias. A lower ENC value indicates having strong ENC bias. The GC3 was calculated as the proportion of guanine and cytosine nucleotides at the third codon position across all coding sequences, reflecting the influence of mutational bias on codon choice. The mean, standard deviation, and quartile for ENC, GC3, and RSCU were summarized to identify a species-level patterns. The variation in ENC and GC3 between the lineages was calculated using one-way ANOVA with the HSD test. The relationship between ENC and GC3 was calculated using Pearson’s correlation coefficient (p < 0.05). A genome-wide pattern of codon usage for ENC and GC3 was visualized in a histogram, while the scatterplot for ENC and GC3 was plotted using kernel density estimation to highlight distributional features. All the analyses were performed using Python-based pipelines.

### GC1,2 Vs GC3 calculation

Genome-wide GC content was calculated for the 1^st^ (GC1), 2^nd^ (GC2), and 3^rd^ (GC3) positions of the codon. The contribution of each codon to the GC content at the specific codon position was determined by assigning values of 1 for G/C and zero for A/T. The species-level GC1, GC2, and GC3 values were calculated as the weighted sum of codon frequencies multiplied by the binary GC contribution, and GC1,2 was calculated as the mean of GC1 and GC2. The neutrality analysis was conducted by fitting a linear regression model of GC1,2 against GC3 for all species. The slope measures the amount of selection pressure relative to the codon-usage constraints. The regression coefficient (slope) and intercept were reported as quantitative measures. The significance of the linear association was studied using Pearson’s correlation between GC3 bases and GC1,2 bases (*p < 0.05*). The relationship was plotted using scatter plots having a fitted regression line, showing species-specific variation and the extent of neutrality. All the analyses were conducted using Python-based pipelines.

### Parity Rule 2 (PR2)-Bias

First of all, the percentage frequency of genome-wide codon usage data was converted to proportions for a more appropriate numeric distribution. The GC content at the 1^st^, 2^nd^, and 3^rd^ positions (GC1, GC2, and GC3) was calculated for each species by weighing the presence of G and C nucleotides at each position by codon frequencies. The mean GC content of the 1^st^ and 2^nd^ positions (GC1,2) was calculated to evaluate the mutational bias relative to the 3^rd^ positions. To assess the parity rule 2 (PR2) bias, the relative position of A nucleotides versus T and G nucleotides versus C at the 3^rd^ position of the codon were quantified as AT-bias [A3/(A3+T3)] and GC-bias [G3/(G3+C3)], respectively ^33^. The missing values were assigned 0.5 to reflect neutrality. The species were classified into the codon bias categories based on their effective number of codons (ENC). The GC3 quartiles were computed to stratify the dataset. The PR2-bias patterns were plotted by GC3 quartiles and ENC across lineages with a neutral expectation of 0.5 for AT and GC bias axes. A linear regression of GC1,2 against GC3 was performed to quantify the influence of mutation pressure, with the regression slope indicating the neutrality. All analyses were done using a Python-based pipeline.

### Machine learning and Correlation Analysis

The codon usage data were used to understand which codons are playing a determined role in codon usage across the lineages. We conducted a decision tree regression analysis in JASP (version 0.95.4.0), using codons as the target variable. The model included 210 training and 52 test sets, with a mean squared error of 1.063e-6. A random forest analysis was performed with a similar number of training and test sets. Each tree used 50% of the training data. The random forest analysis provided node purity, mean decrease in accuracy, and predictive performance of the codons. A neural network analysis was conducted using the JASP software (version 0.95.4.0). A 20% of the test set and 20% of the training and validation sets were used for the neural network analysis. A logistic sigmoid activation function with the Rprop heuristic artificial neural network was used for this ^34,35^. The neural network analysis was optimized for 10 generations, with a maximum of 10 layers and a maximum of 10 nodes in each layer. Pearson’s correlation analysis was conducted to generate a heatmap with 95% confidence to understand the correlation between the codons using the JASP software (version 0.95.4.0). We used a weighted network plot with a non-parametric bootstrapping to generate the codon network in JASP (version 0.95.4.0). Principal component analysis of the codon usage with their lineages was conducted using Unscrambler software. Clustering (Pearson’s) of codons based on their usage and a bar graph of the codon frequency were generated using Past4 software.

## Results

### Comparative codon analysis showed non-random and AT-based codon usage and nucleotide preference across plant genomes

Genome-wide codon usage analysis of about 4.6 billion (4,617,221,004) codons of 11,372,724 CDS sequences from 262 plant species. It constituted approximately 43407 CDS and 17622980 codons per species. In the study, *Triticum dicoccoides* contained the highest number (295,286), while *Cyanidioschyzon merolae* contained the lowest number (252) of CDS (**Supplementary Table 1**). The highest percentage of codons found during the study was GCG (8.42%) in *Raphidocelis subcapitata* followed by GGC (6.66%) in *Chlamydomonas schloesseri*. The lowest percentage of codons was TTA encoded by *Gonium pectoral* (0.054%, excluding the stop codons). A few codons, GAA (*Coffea arabica*), GAT (*Coffea arabica*), and GGC (*Chlamydomonas reinhardtii*) were found to be absent in the CDS of their entire genomes (**Supplementary Table 2**).

Analysis of synonymous codon usage revealed a non-random bias in the codon choice which varies among amino acids but showed a genome-wide trend. Usage is unevenly distributed, indicating a combination of mutational bias and selective constraints rather than neutral usage. Amino acids Arg, Leu, and Ser, encoded by six synonymous codons provide synonymous codon usage asymmetry. For example, for the Arg amino acid, the codons AGA and AGG (codons ending with A or G) show higher average usage compared to the GC/CG-rich codons (CGT, CGC, and CGG) (**Supplementary Figure 1**). This suggests avoidance of CpG-containing codons that are associated with mutational pressure. The amino acid Leu preferred TTG, GTT, and CTT, whereas CTA is under-represented (**Supplementary Figure 1**). Similarly, for Ser TCT, TCA, and AGC were preferred, while TCG was less preferred, reflecting the suppression of GC-rich codons. In the four-codon family (Ala, Gly, Pro, Thr, and Val), the synonymous codon usage is biased towards the codon ending with T or A. In Ala, codons GCT and GCA; in Gly, GGT and GGA; in Pro, CCT, and CCA; in Thr, ACT and ACA were dominated over their other synonymous codons encoding with a C or G nucleotide (**Supplementary Figure 1**). Val showed a strong preference for GTT and GTG, while GTA was not favored. These patterns strongly indicated a genome-wide bias towards favoring A/T-encoding codons in plants. The two codons encoding amino acids (Asn, Asp, Cys, Gln, Glu, His, Lys, Phe, and Tyr) showed synonymous bias (**Supplementary Figure 1**). In most cases, the codons ending with T (AAT, GAT, TTT, TAT) were more abundant, with an exception to Lys, where AAG was favored over AAA, reflecting amino acid-specific deviation from A/T-rich codon preference. The single-codon amino acids Met (ATG) and Trp (TGG) showed no bias, while the stop codon TGA was more favored than TAA and TAG. This shows a preferential termination signal across the plant genome that might be linked to translational efficiency. Genome-wide nucleotide position analysis revealed distinct positional bias. Adenine shows the highest frequency at the 2^nd^ position of the codon, followed by the 1^st^, and 3^rd^, while guanine has the highest representation at the 1^st^ position, followed by the 2^nd^ and 3^rd^ positions (**Supplementary Figure 2**). Thiamine nucleotide is preferred at the 3^rd^ position followed by the 2^nd^ and 1^st^ positions, while cytosine nucleotide is preferred at the 2^nd^ position, followed by the 3^rd^ and 2^nd^ positions (**Supplementary Figure 2**).

Parity rule 2 (PR2) analysis revealed that there is a clear departure from the expected equilibrium for the complementary nucleotide at the 3^rd^ position of the codon. The GC-bias (G3/[G3+C3]) and AT-bias (A3/[A3+T3]) values were distributed quite unevenly around the central 0.5 axis (Figure 1A). This indicates mutation and selection events do not act symmetrically on the fourfold degenerate sites. Most species clustered towards an A- and G-rich quadrant. This shows the preferential accumulation of purines at the synonymous positions. In lineage-specific analyses, monocots and eudicots occupied partially overlapping but directionally skewed distributions. Eudicots tended towards the higher AT asymmetry, while monocots showed comparatively greater skew. Bryophytes and algae showed dispersed distributions, reflecting heterogeneous mutational pressure in the early-diverging plant groups. Pteridophytes and basal angiosperms clustered towards the center, suggesting weaker PR2 deviations (Figure 1B). This lineage-specific displacement highlights the role of clade-dependent genome composition, selective constraints, and replication bias on the evolution of synonymous sites. The quartile-based stratification also shows internal PR2 variations (Figure 1C). Each quartile shows a bias pattern. The species in the GC-quartile showed GC skew with limited AT asymmetry, while the species in the lower GC quartile showed a more variable A3/T3 ratio and higher dispersion near the axis (Figure 1C). The spread across the quartile suggests that PR2 deviation rose because of a combination of mutation-selection balance and the lineage-specific genomic landscape, rather than uniform, stochastic fluctuations. This shows PR2 bias is a robust indicator of asymmetric nucleotide usage across the plant kingdom, and the deviation from parity varies with evolutionary lineage and genomic backgrounds.

**Figure 1.**
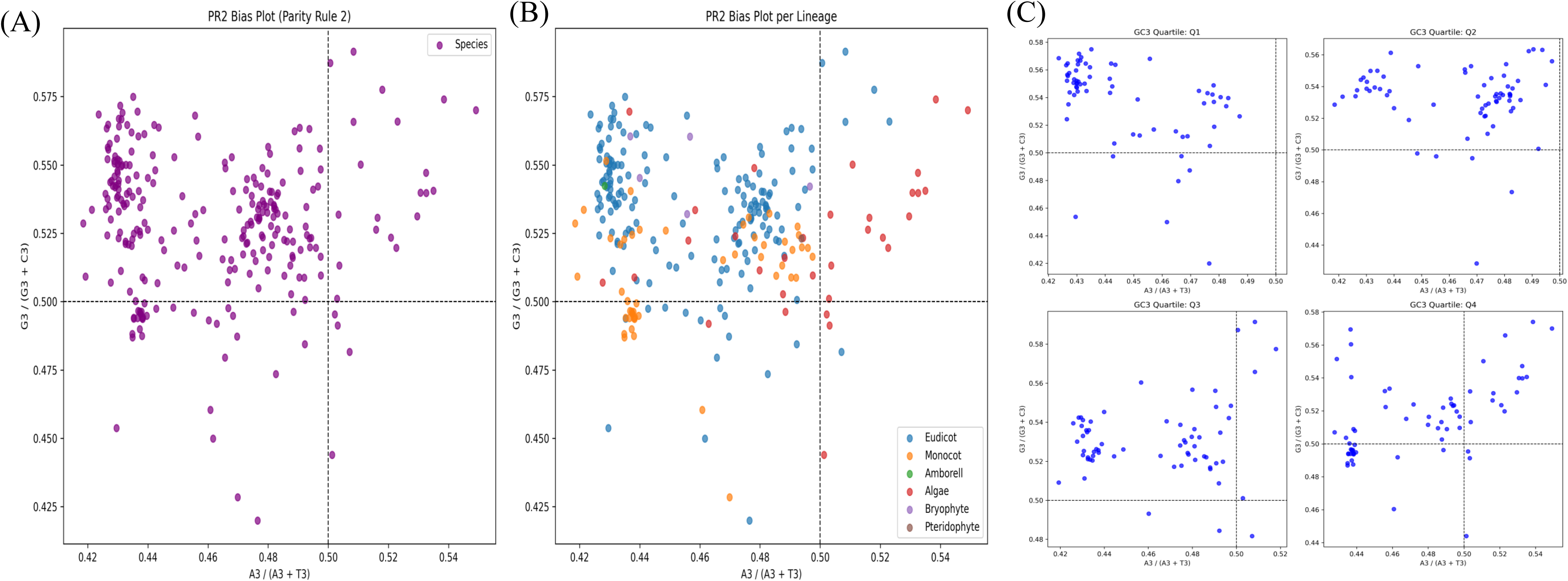
PR2-bias analysis across species, lineages, and GC3 content strata. (A) Parity rule 2 (PR2) bias plot showing the distribution of [A3/(A3+T3)] versus [G3/(G3+C3)] across all species analyzed. Each point represents one species, showing genome-wide deviations from PR2 expectations at the third codon position. The horizontal and vertical dashed lines denote the PR2 equilibrium at 0.5 for AT and GC, respectively. (B) Represents the PR2-bias plot partitioned by major evolutionary lineages, indicating eudicots, monocots, algae, bryophytes, and pteridophytes. Distinct lineage-specific clustering patterns highlight variation in asymmetries. Figure (C) represents PR2-bias plots that are stratified by GC3 quartiles, as seen in Q1-Q4. The species are binned according to increasing GC3 abundance to understand the relationship between GC compositional constraints and PR2 deviation. GC3 enrichment is associated with systematic shifts in AT and GC imbalance in all the quartiles, indicating differential mutation and/or selective constraints acting on synonymous sites.

### Genome-wide codon and amino acid usage bias shows translational selection and lineage-specific evolutionary constraints

Based on the codon frequency, we plotted the corresponding amino acids. The analysis showed that Leu encodes the most codons, contributing about one-tenth of the amino acid pool, followed by Ser, Arg, Ala, and Gly (Figure 2A). The codons for these amino acids dominate the genome, indicating a strong bias towards highly degenerate codon families. In contrast, Met and Trp, which are encoded by only one codon, showed the lowest frequencies, while stop codons are present in minimal amounts (Figure 2A). The gradual decline in codon usage across amino acids reflects codon degeneracy and the selective constraints on protein composition and translational efficiency. The study also shows heterogeneity in synonymous codons, which strongly suggests codon usage bias. The frequency of the AAG codon was the highest, followed by GAA, GAG, GAT, and AAA (Figure 2B). Some codons occurred more frequently than their synonymous counterparts. This suggests that preferential codon usage is most possibly driven by tRNA availability and/or translational optimization (Figure 2B). The codons whose nucleotides ended with G or C showed higher usage in a few cases, suggesting the role of GC content in codon preference. The PCA plot shows that synonymous codons clustered quite closely, suggesting their role in shared compositional and functional constraints (Figure 1C). In PC1, the codons with high GC percentage are separated from the AT-rich nucleotides, thus underscoring the important role of nucleotide composition in shaping the codon usage pattern (Figure 2C). PC2 shows lineage-specific grouping of codons where algae are distinctly separated while monocots, eudicots, Magnoliids, and Nymphaeales partially overlap (Figure 2C). This pattern suggests lineage-specific codon usage, reflecting evolutionary divergence and differential selective pressures across plant lineages. The proximity of higher plants shows conserved codon usage strategies, while separated algae highlight a shift in translational and genomic architecture during the evolution of land plants.

**Figure 2.**
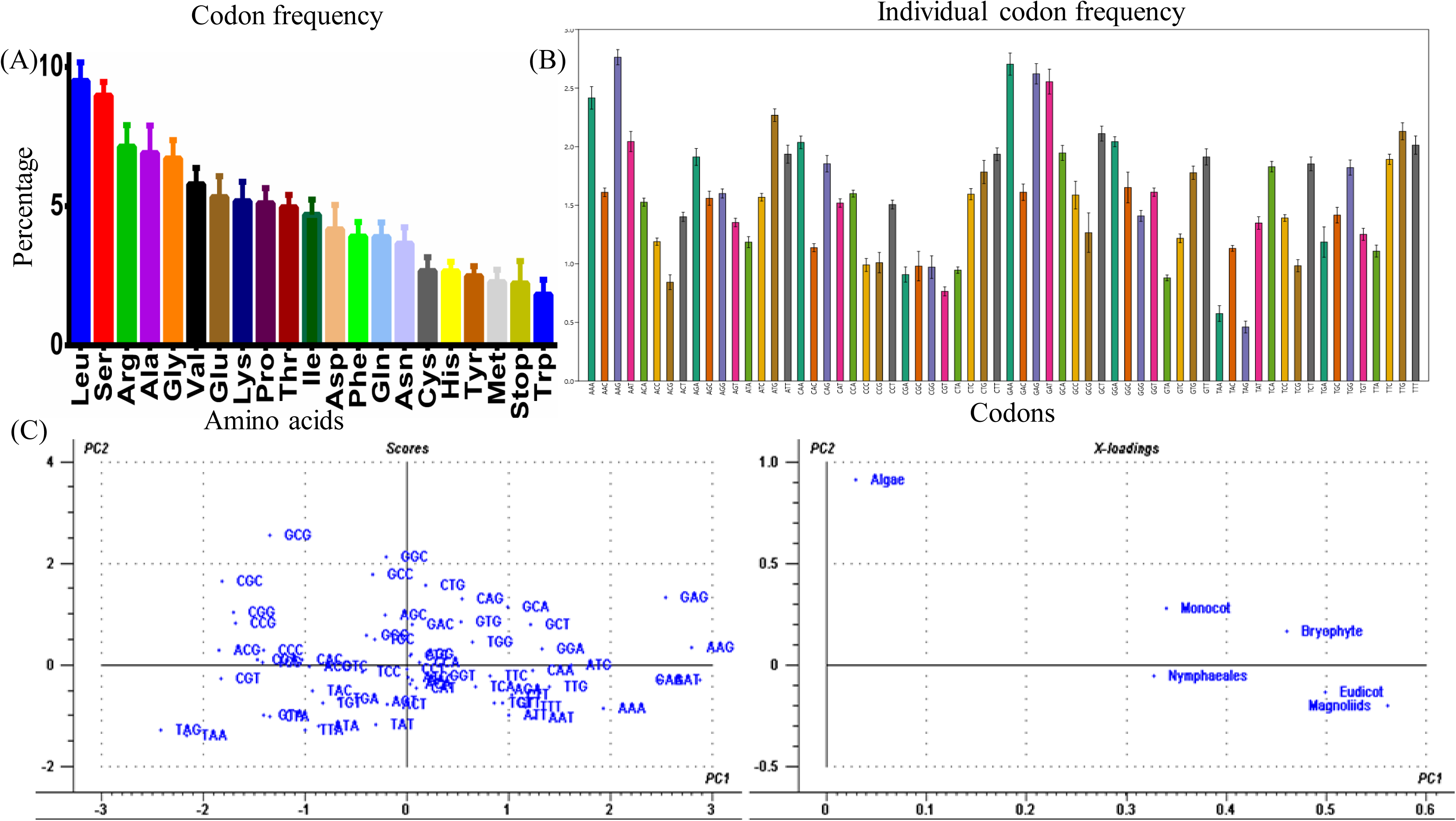
Codon usage pattern, codon-specific frequencies, and multivariate analysis of codon usage bias. Panel (A) represents overall codon usage frequencies at the amino acid level. The bar chart shows the cumulative proportion of synonymous codons used for each amino acid in increasing to decreasing order. The error bars indicate variability among species, capturing differences arising from selective mutational pressure that shaped amino acid composition. Panel (B) represents the distributions of individual codon frequencies with error bars. This reveals considerable variation in synonymous codon usage, with some codons overrepresented and others underrepresented. Panel (C) depicts the principal component analysis of the codon usage patterns. The left plot shows the distributions of the selected codons. PC1 and PC2 represent the primary axes explaining variance in synonymous codon usage bias. The right panel shows PCA loadings. This reveals clustering of major evolutionary lineages and emphasizes lineage-specific patterns of codon usage. Overall, the figure indicates that the codon usage bias is conserved, consistent, and phylogenetically structured across the plant lineages.

### Codon adaptation index (CAI) showed genome-wide optimization of translational efficiency

The codon adaptation index (CAI) of the codon usage was studied by comparing the codons of the CDS with the highly expressed genes of *Arabidopsis thaliana*. The study showed that the CAI of *Cucurbita maxima* were highest (0.77), whereas the CAI of *Gonium pectorale* was lowest (0.57) (**Supplementary File 1**). The results showed a non-uniform distribution of CAI values and a pronounced skew towards the higher values. The dominant peak is centered around 0.72-0.75 (Figure 3A). This indicates that most of the genes showed strong adaptation towards preferred codons, consistent with accuracy and translational efficiency. The tail extending toward lower CAI values (∼0.57-0.65) suggests genes or gene pools are under relaxed selection or lineage-specific regulatory constraints. Although the overall distribution is heterogenous, codon optimization is predominantly high across the species. This highlights the role of natural selection in shaping the codon usage pattern. Figure 3B shows the preferred codon usage for each amino acid. The results show a clear dominance of synonymous codons within amino acid families, indicating a clear role of codon usage bias. Gln, Arg, and Glu showed highly preferred codon usage at 3% or more in some cases (Figure 3B). This indicates a strong selection for optimal codon usages. However, the amino acids with lower degeneracy and reduced functional demands exhibited modest preferred codon frequencies. The majority of preferred codons have GC-endings, showing the role of nucleotide composition in codon optimization (Figure 3B). The concordance between the high CAI values and preferred codon frequencies supports CAI as an indicator of translational selection and adaptive codon usage. The dendrogram groups codons into three major clusters (Figure 3C). These clusters are well defined and correspond to their synonymous relationships and compositional properties (Figure 3C). Codons coding for the same amino acids tend to cluster together, showing their coordinated usage pattern and shared evolutionary pressures. The GC-rich codons formed distinct clusters separated from AT-rich codons, reflecting the role of nucleotide composition as a major determinant of codon grouping (Figure 3C). The codons TCT, TTT, TGT, CTT, and CCA, which are highly expressed in the genes, are clustered together (Figure 3C). Clustering of codons close to each other with similar usage frequencies reflects coadaptation of codons potentially driven by tRNA abundance and translational efficiency. The CAI-based study (Figures 3A and 3B) and the codon-based clustering (Figure 3C) further support strong and non-random codon usage bias. Compositional and functional constraints governed the alignment of high CAI values with higher usage of particular preferred codons and coherent codon groups. These findings show that codon usage patterns are shaped by an intricate interplay among nucleotide composition, translational selection, and evolutionary history. This suggests a mechanistic insight into the genome-wide optimization and gene expression.

**Figure 3.**
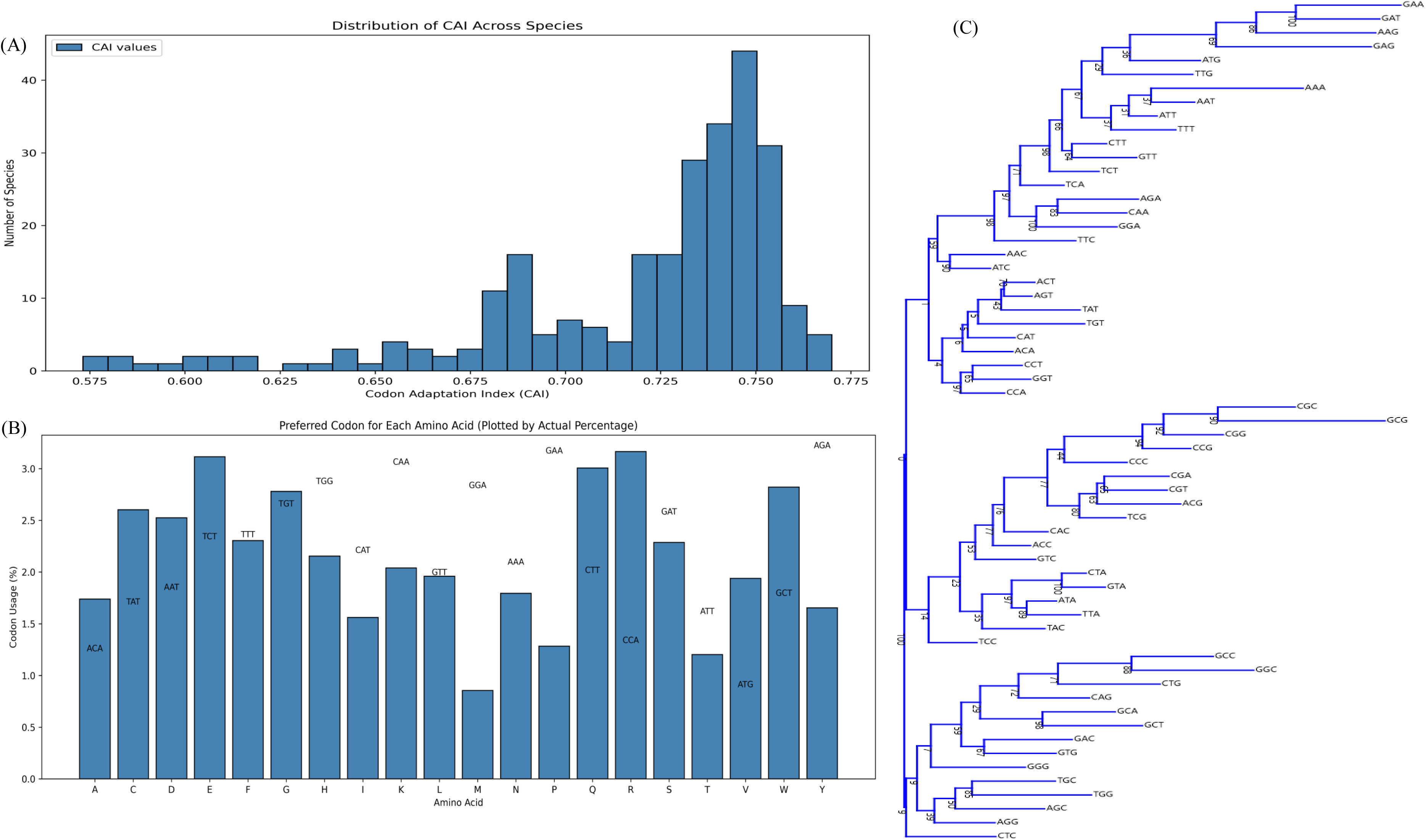
Codon adaptation-index (CAI), preferred codon usage, and codon-based phylogenetic relationship in the plant kingdom. Panel (A) represents the distribution of CAI values, while the histogram represents the notable variation in the CAI. Most species cluster toward higher values, reflecting a general trend of codon optimization relative to highly expressed genes. These differences highlight a considerable variation in translational efficiency and the strength of codon selection across the species in the lineage. Panel (B) represents the preferred codon usage for each amino acid, expressed as a percentage of total synonymous codons. For each amino acid, the dominant codon is plotted, showing a clear preference for specific synonymous codons. This reflects that selection is driven by the translational efficiency and tRNA abundance in the genome. Panel (C) represents a codon-based phylogenetic tree that group synonymous codons by shared substitution patterns and compositional similarities. The clusters reflect constraints on codon usage shaped by evolutionary forces. The branching structure represents the organization of codon preferences and their divergence over evolutionary time.

### Lineage-dependent codon usage optimization is driven by codon-anticodon co-evolution in plants

Analysis of preferred codons with relative synonymous codon usage (RSCU) greater than one across the plant lineage showed an ordered stratification rather than a random distribution (Figure 4A). They exhibited a broadly conserved yet lineage-modulated codon usage. The monocots displayed the highest number of preferred codons, showing a relatively stronger codon usage bias, whereas the eudicots displayed a slightly reduced but robust preference for codons. The arboreal and algae showed comparatively reduced bias, indicating a weaker and heterogeneous codon bias. The Bryophytes and Pteridophyte showed intermediate bias, suggesting a gradual evolutionary transition in codon optimization from basal to diverged lineages. Although lineage-specific differences are evident, a few codons are shared across groups. This supports the retention of a conserved translational pool that is differentially redefined during plant evolution. Further, codon-anticodons pairs were mapped using percentage-frequency metrics, which showed a highly structured landscape of translational coupling (Figure 4B). Hierarchical clustering segregated the codons into discrete modules corresponding to the anticodon abundance. This shows a non-random codon-tRNA association (Figure 4B). High-frequency codon-anticodon pairs cluster tightly, reflecting selective enrichment of cognate and functionally efficient near-cognate interactions. For example, GAA (Glu) and AAA (Lys) codons showed high frequency and formed tight clusters with their cognate anticodons UUC and UUU. Respectively, indicating strong translational demand supported by tRNA abundance. Similarly, GCU/GCC (Ala) and CCU/CCC (Pro) clusters with their respective anticodons, reflecting preferential use of GC-rich synonymous codons in lineages with expanded tRNA pools for these amino acids. In contrast, the low-frequency codon-anticodon pairings remain diffuse, consistent with the limited translational demand or relaxed selective pressure. For example, AUA (Ile) and UUA (Leu) showed a lower frequency and weaker clustering, which is consistent with limited anticodon abundance and reduced translational optimization. A near-cognate interaction was observed within the clusters of synonymous codons (Figure 3B). The codons UCU, UCC, AGU (Ser) and CGU, CGC, AGA (Arg) showed functional flexibility (Figure 3B), allowing efficient decoding despite the Wobble-base pairing. This showed a coordinated adjustment between codon usage and cellular tRNA supply rather than stochastic usage of synonymous codons. These codon-anticodon pairings demonstrated that codon usage is not merely associated with its synonymous redundancy, but reflects a finely orchestrated co-evolution between codon preference and anticodon availability, optimizing translation efficiency in a lineage-dependent manner.

**Figure 4.**
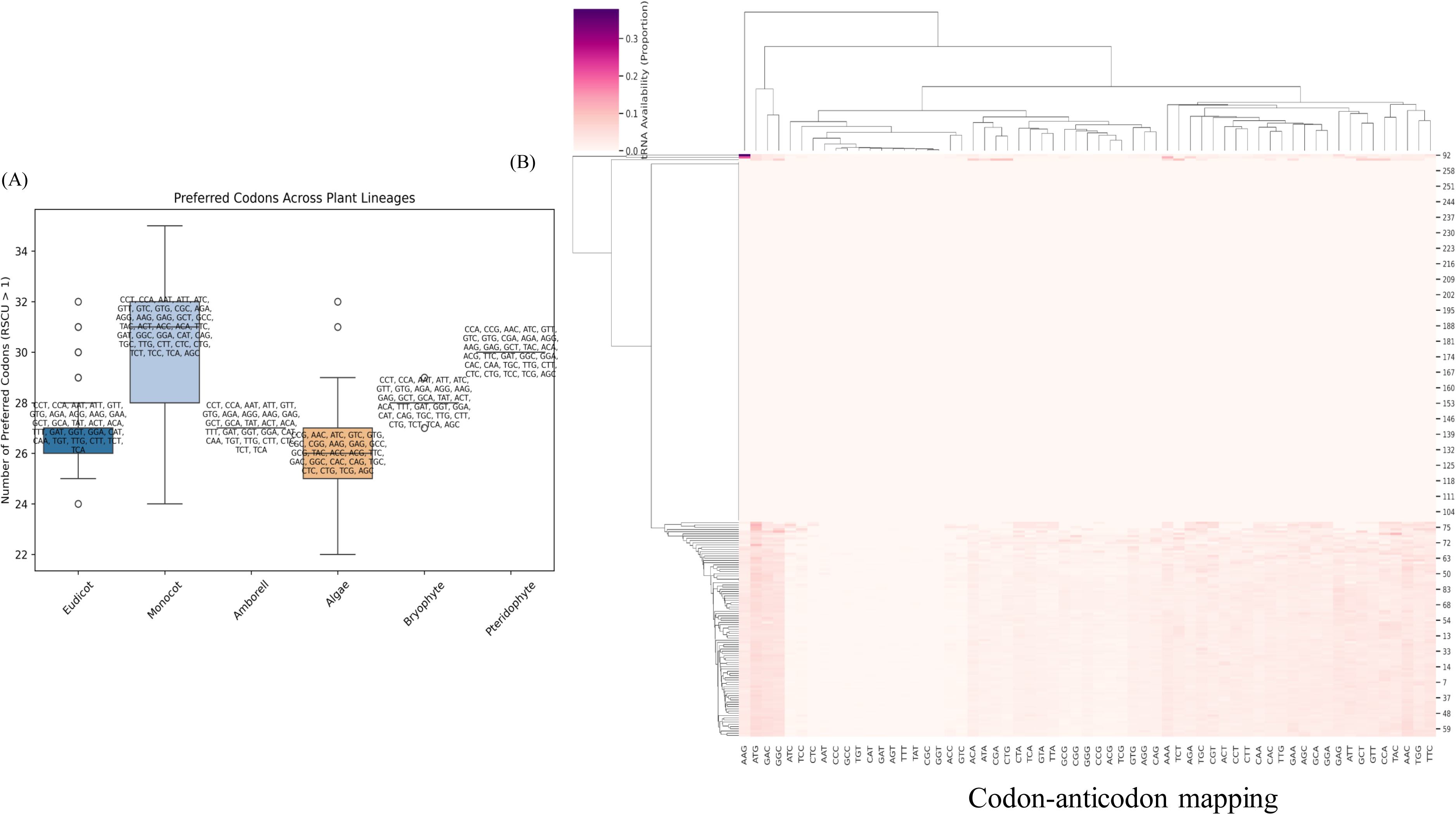
Lineage-specific codon usage and codon-anticodon relationship across the plant lineages. Panel (A) represents the distribution of preferred codons with RSCU > 1 across the major plant lineages. The box plot summarizes the number of preferred codons identified within different lineages in the plant kingdom. The preferred codons were annotated for each lineage, reflecting lineage-dependent biases. This highlights the preference for conserved versus diverged codon patterns across the plant kingdom’s evolutionary lineages. Panel (B) represents the hierarchical clustering heatmap of codon-anticodon mapping. Color intensity represents relative codon-anticodon usage frequency, with darker shades indicating stronger bias. The dendrogram shows codon and taxon clustering, reflecting coherent modules of codon-anticodon co-adaptation. Overall, these patterns indicate coordinated evolution of codon-usage preferences and tRNA anticodon pool across diverse plant lineages. Basal lineages show relatively diffuse, less specialized codon preferences, whereas diverged lineages like monocots show increased codon specialization. This strong concordance between codon and anticodon frequencies indicates codon-anticodon co-evolution via translational selection. This highlights lineage-specific codon optimization of translational efficiency while maintaining the conserved structure of the genetic code.

### Decision tree and neural network analysis revealed CTC codon as the primary determinant and a major predictor

A machine learning strategy was implemented to determine the codons with the highest predictive contributions using complementary tree-based and neural network models. The decision tree regression analysis showed that the CTC codon emerged as the primary split variable at the root node (Figure 5A). This reflects that the CTC codon account for a considerable portion of the variance in the response variable. This emphasizes CTC as the dominant codon influencing all the other codons. The subsequent branches provided only minor refinements. The terminal nodes distinguished only a subset of observations characterized by the extreme values. The lowest mean square error indicated that the model achieved a strong predictive performance without overfitting (Figure 5A). This finding supported the biological significance of a limited number of informative codons rather than genome-wide distributions.

**Figure 5.**
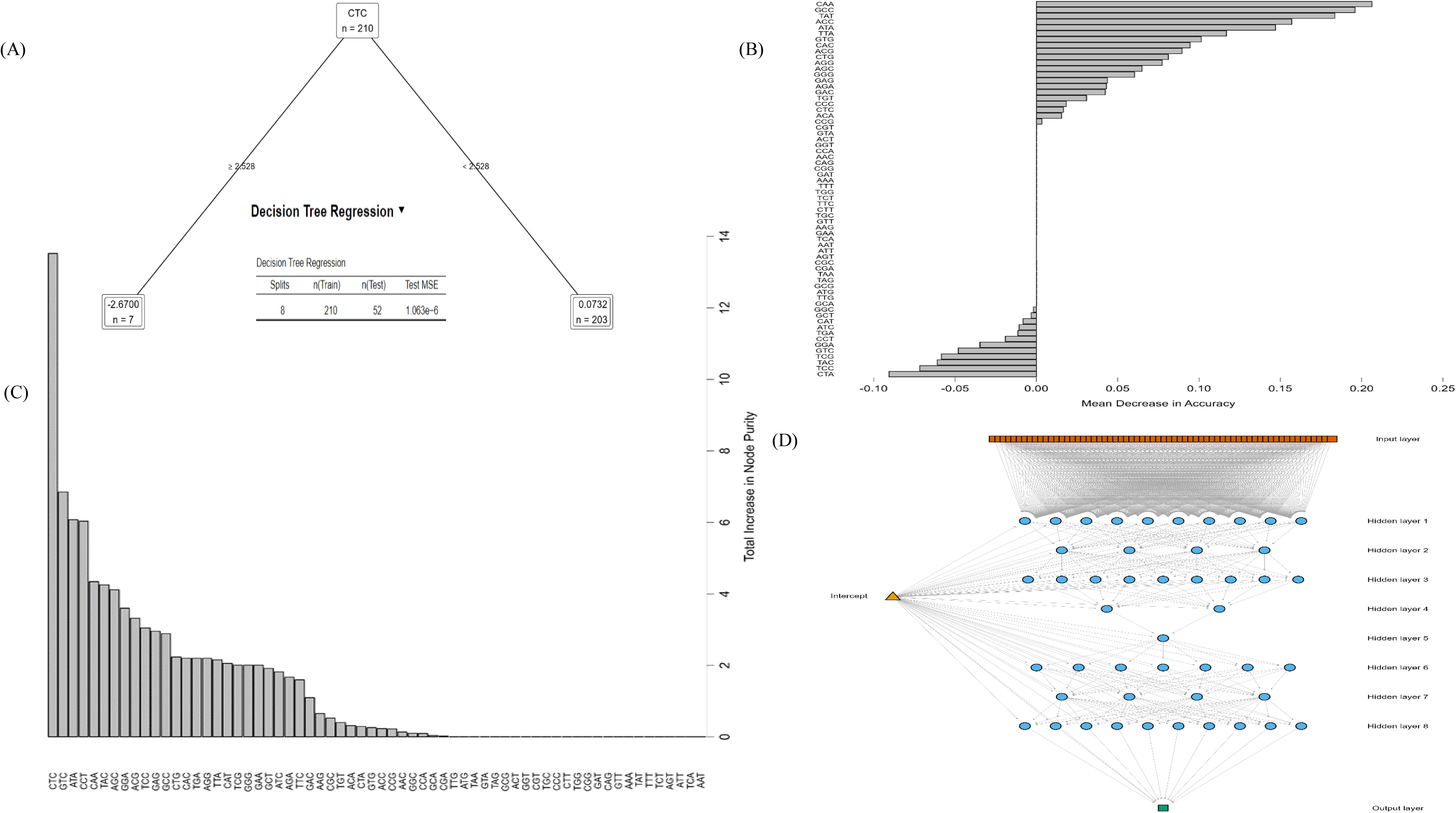
Machine-learning-based identification of influential codons and predictive modeling of codon-associated features. Figure (A) represents decision tree regression that identified key codons contributing to the predictive model. The tree illustrates hierarchical partitioning of the dataset (n = 210), where codon features are recursively split to minimize prediction error. The terminal nodes represent subsets of observations with similar predicted values, and the model summary (inset) reports the number of splits, training, and test sample sizes, and mean squared error (MSE). (B) Variable importance estimated using the mean decrease in accuracy. The codons are ranked by the reduction in model accuracy when each feature is permuted, indicating their relative contribution to the predictive performance. The positive values indicate codons whose perturbation strongly decreases model accuracy, whereas near-zero or negative values denote features with minimal predictive contribution. (C) Variable importance is measured as the total increase in node purity across the model. (D) represents the architecture of the artificial neural network used for predictive modeling. The network contains an input layer representing the codon features, multiple fully connected hidden layers, and single output node. The single node generates the final prediction. The weighted connections between the layers provide nonlinear interactions with the codons. This enables the model to learn complex hierarchical relationships in the codon usage patterns.

The mean decrease in accuracy (MDA) of random forest analysis identified CAA as the most influential codon, followed by GCC, TAT, ACC, and others. Several codons clustered near zero, reflecting their minimal contribution to the predictive performance. A few codons, including CTA, TCC, TAC, TCG and others, showed negative MDA values. This suggests redundancy or negligible predictive influence. This uneven distribution supports a model where evolutionary and translational signals are concentrated in particular synonymous codons rather than distributed uniformly across the genome (Figure 5B). Figure 5C reflects that similar codon, those ranked highly by accuracy, also produce the greatest cumulative reduction in node impurity. This shows consistent, robust splitting behaviour across the ensemble. This indicates a stable association with the modelled biological outcome. Ranking based on node purity and accuracy further emphasize the robustness of these codons as key explanatory variables.

The neural-network analysis (Figure 5D) showed a highly interconnected architecture with numerous hidden layers linking the codon-level inputs to outputs. This suggests a complex and non-linear connections. The model captured synergistic and complementary interactions that impact genome-wide codon usage by integrating dispersed signals across multiple codons. The codons identified in tree-based analysis also showed a stronger connectivity pattern within the network, suggesting they serve as the central hubs in learned representations. This convergence across the modelling paradigms indicates that both non-linear combinatorial effects and linear separability contribute to the codon-level predictive structure.

### Genome-wide codon usage bias driven by GC3-associated mutational pressure

The genome-level distribution of effective number of codons (ENC) exhibits lineage-dependent stratification (Figure 6A). Eudicot genome clusters predominantly in a high ENC-area with a bimodal distribution, centered around the mid-to-high ENC range. This shows a relatively weak codon usage bias across multiple species. However, the subgroups show a stronger constraint. The monocot species showed a partially overlapping but right-shifted distribution, indicating moderate codon bias but with a narrower dispersion than the eudicots. In contrast, the early-diverging lineage algae showed lower ENC values. The algae species showed the widest dispersion and the lowest ENC range, consistent with strong codon usage bias and extreme compositional constraints. Bryophytes and pteridophytes occupied intermediate positions, with reduced ENC compared to the angiosperms. However, ENC was higher than that of algae, suggesting a gradual relaxation of codon bias during plant evolution. The presence of a single *Amborella* species towards the lower end of the angiosperm ENC, reflects its transitional evolutionary status between basal angiosperms and flowering plants. The distribution shows that codon usage bias is not uniform across lineages, and the genome-wide codon usage freedom increased progressively from algae to the higher angiosperms.

**Figure 6.**
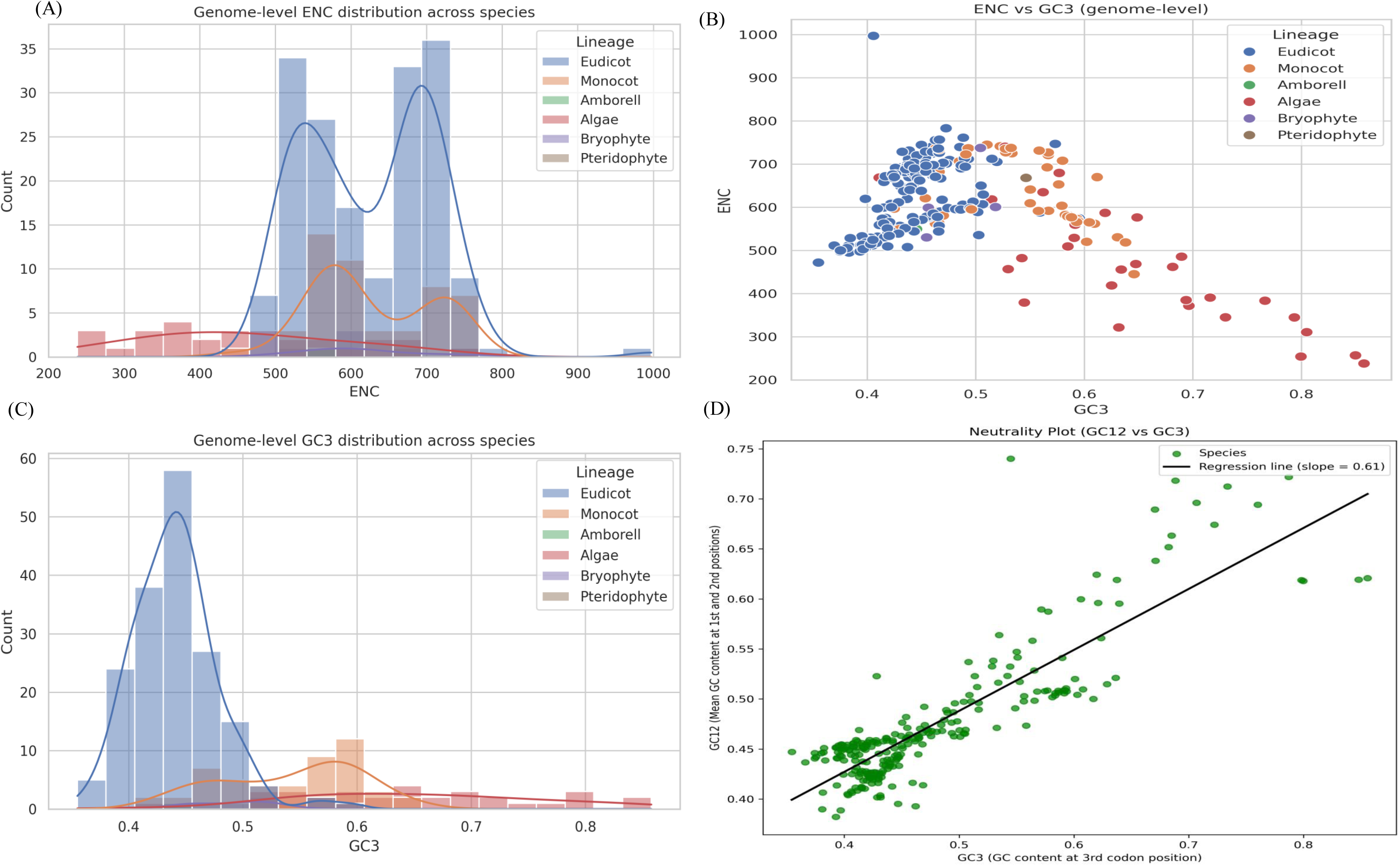
Genome-wide patterns of codon usage bias and GC composition across plant lineages. Panel (A) shows distributions of effective number of codons (ENC), and the histogram with kernel density overlays illustrates lineage-specific variation in codon usage bias. The high codon usage values reflect more uniform codon usage values reflect more uniform codon usage, whereas the lower ENC indicates stronger codon usage bias. Panel (B) shows the relationship between ENC and GC content at the third codon position (GC3). Each point in the figure represents a species. The distribution shows lineage-specific clustering. This suggests that variation in GC3 is associated with differences in codon usage bias across plant genomes. Panel (C) represents the genome-level distribution of GC3 content across the lineages. This histogram with the density curve shows variation in GC3 content across key plant lineages. This indicates variability in nucleotide composition and possible lineage-specific mutational or selection pressure affecting the codon usage. Panel (D) show the neutrality plot evaluating the correlation between GC content at the first and second positions (GC1,2) and at the 3^rd^ position (GC3). The regression line has slope of 0.61, indicating that mutational pressure shapes codon composition and that codon selection affects nucleotide composition at coding sites.

The scatter plot shows the genome-level ENC versus GC content at the third position. It shows an inverse relationship across all the examined lineages, where genomes with elevated GC3 values consistently exhibited reduced ENC (Figure 6B). This indicates strong codon-usage bias under the GC-rich conditions. The algae lineage occupied the extreme high-GC3 and low-ENC quadrant, showing strong compositional pressure at the synonymous sites. The eudicots and monocots formed partially overlapping but clearly visible boundaries (Figure 6B). Monocots extended towards higher GC3 and lower ENC values than the eudicots. This suggests strong GC-driven-codon constraints in monocot genomes. Eudicots were dominantly concentrated in a moderate GC3 range with high ENC (Figure 6B), reflecting a weaker compositional bias and greater codon flexibility. However, the ENC-GC3 trend showed no lineage-specific patterns and supported universal composition-driven constraints acting on synonymous codon choice at the genome scale. As depicted in Figure 6C, eudicots showed low GC3 distribution, whereas monocots displayed a clear shift towards the GC3 values with broader variance. The algae lineage showed extreme GC3 enrichment across a wide range, consistent with their low ENC and strong codon bias. Bryophytes and pteridophytes occupied the intermediate position, suggesting the modulation of GC3 compositional force during adaptation and evolution of terrestrial vascular plants. This explains that GC3 is the primary axis of evolutionary divergence in the plant genome and is tightly related to codon usage bias. Figure 6D explains the neutrality plot that compares the GC content at the first and second codon positions (GC1,2) against GC3. It shows a strong positive correlation with a fitted regression line showing a slope of approximately 0.61. This indicates that changes in GC3 explain about 61% of the variation in GC1,2. The slope provides evidence that mutational bias is the dominant force shaping codon composition, but not the sole determinant. The remaining unexplained variance suggests some selective constraints, such as nonsynonymous positions modulating the base composition. This shows that the balance between mutation and selection operates in a quantitatively similar manner across the lineage despite the differences in ENC and GC3 values.

### Deep evolutionary divergence in plants revealed by distinct codon-usage signatures

A global codon usage profile revealed compositional and preferential differences between the higher and lower plant groups (Figure 7A). The heatmap showed pronounced differences in relative synonymous codon usage (RSCU) across two clades. Higher plants contained A/T-ending codons (AAA, AGA, TTA, TTT), while the lower plants showed comparatively elevated G/C-ending codons (CCG, GGC, GAG) (Figure 7A). This reflected the GC-biased mutational context and ancestral genome architecture of the algae and Bryophyte lineage. These patterns reflect the transition from lower to higher lineages, accompanied by a shift in codon-usage bias, most likely driven by changes in the genome composition and lineage-specific selective pressure acting on protein translation machinery efficiency. Figure 7B shows lineage-level heterogeneity, where the algae lineage shows a GC-rich signature while monocots show distinctive enrichment towards A-or-G ending. This is consistent with their known bias in genomic GC content and tRNA abundance. Eudicots showed an intermediate usage pattern, with clade-restricted enrichment of AGG, GGA, and TCA. This result showed the influence of phylogenetic history, mutational dynamics, and translational optimization in shaping codon preference in the evolution of plant lineages. Figure 7B showed further divergence in codon-usage optimization among major lineages. The angiosperms showed a strong selection for reduced preferred codons, while monocots favored C-ending codons. The eudicots showed a higher level of A/T-ending codons such as ATT, TTA, and TTT (Figure 7C). Bryophytes and algae prefer GC-rich codons, clearly separating them from angiosperm profiles. However, the CTA codon is highly enriched in the monocot lineage relative to others. The results show that codon usage bias in highly expressed genes is not uniform, reflecting the evolutionary divergence in translation optimization strategies.

**Figure 7.**
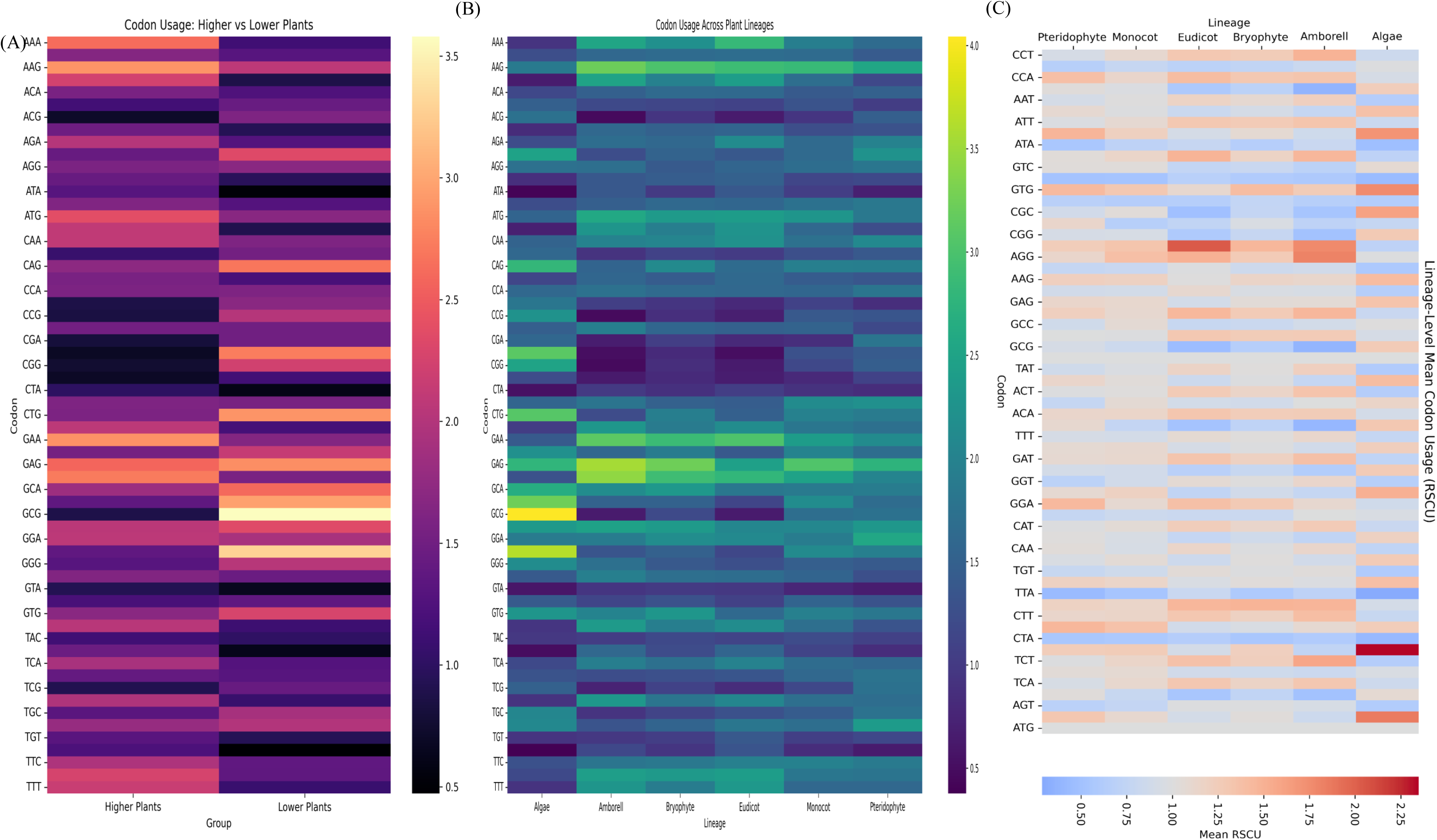
Comparative codon usage trends among major plant lineages. The shows a comparative analysis of relative synonymous codon usage (RSCU) in higher and lower plants. The color intensity reflects the magnitude of RSCU values. This highlights variation in codon choice between the evolutionary groups. The rows represent the codons, whereas the column represents lineage-level averages, reflecting their lineage-specific codon usage patterns. The relative increase and decrease of codons across multiple lineages explain the evolutionary differences in synonymous codon preferences in different plant lineages.

### Molecular economy of codon usage is conserved across the plant lineage

To understand the energetic optimization of codons across the plant kingdom, we calculated the molecular economy of codons. Molecular economy is defined as the ATP expenditure to code a codon. Analysis revealed that *Kingdonia uniflora* has the highest (31.14) and *Manihot esculenta* has the lowest (28.51) molecular economy (**Supplementary File 2**). The mean molecular economy of codon usage in plant genomes was 30.30. The distribution of molecular economy values was multimodal, with most species falling between 29.5 and 31.0 ATP per codon (Figure 8A). A prominent peak at approximately 31 ATP per codon shows that a large set of species needs a higher translational energy requirement, whereas peaks around 29.5 and approximately 30.0 ATP per codon indicate the presence of lineage-specific subgroups. The comparative study showed that the molecular economy values are similar, with a modest difference between lineages (Figure 8B). Most groups clustered around 30 ATP per codon, with a minimal standard deviation. This shows translational energetic costs are highly conserved across evolutionary lineages despite divergent genome architecture and codon-usage patterns. This indicates a strong evolutionary constraint on the energetic efficiency of protein synthesis, with only a minor lineage-specific difference.

**Figure 8.**
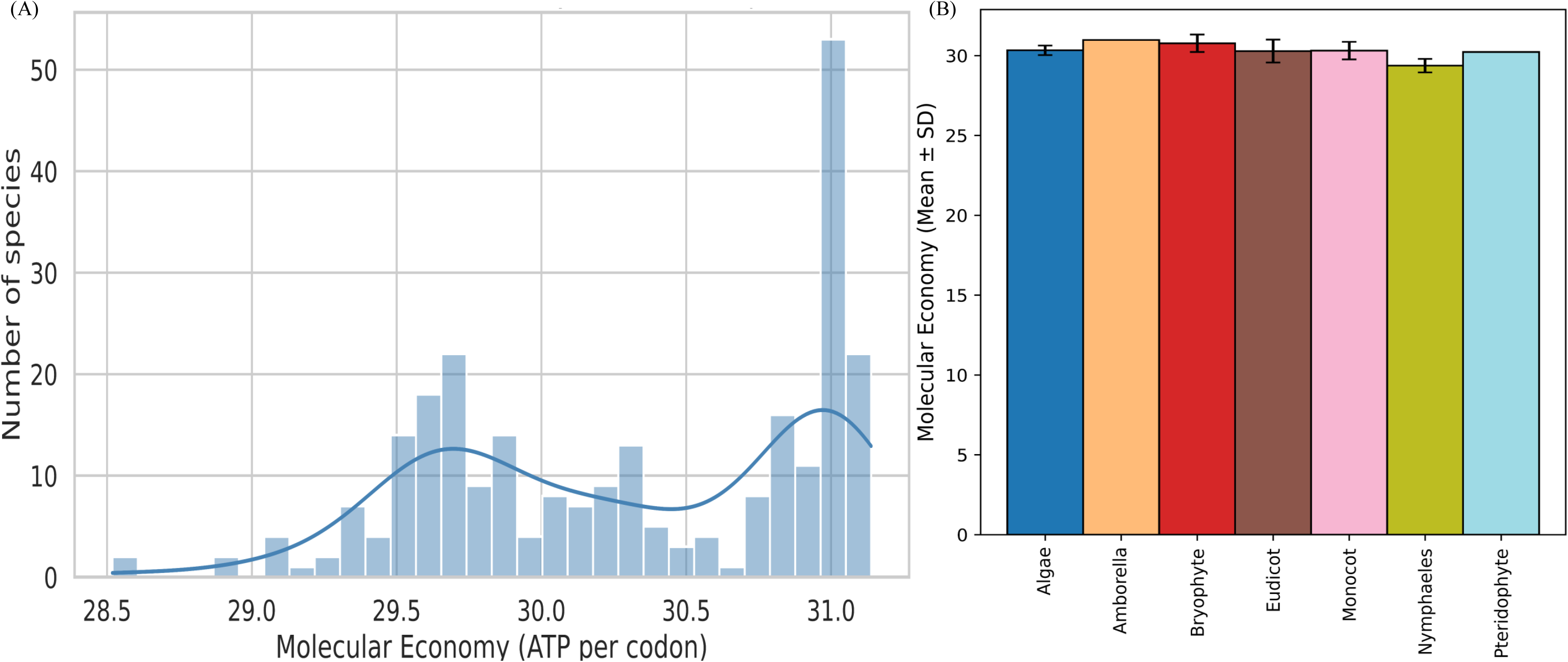
Molecular economy of codon usage across plant lineages. Panel (A) represents the distribution of molecular economy (ATP cost per codon) across all the surveyed species. The histogram bar reflects the species count in each molecular economy bin. The overlaid curve shows a kernel-smoothed density estimate that highlights multimodal patterns in ATP expenditure. Panel (B) explains the mean molecular economy (± SD) in major evolutionary lineages of plants. The bar represents lineage-level mean ATP cost per codon.

### Lineage-specific codon usage correlation and codon usage network show coordinated evolutionary constraints

Correlation analysis across lineages revealed distinct lineage-specific patterns in codon usage relationships (Figure 9A). Algae, Bryophytes, eudicots, and monocots showed different correlation structures across most lineages. The eudicots showed a negative correlation (−0.941) relative to Bryophytes, whereas the monocot showed a strong correlation (0.993). Bryophytes and algae showed a diffuse distribution, indicating more heterogeneous codon-usage behavior within the early-diverging lineages. The dispersed correlation coefficients reflect the evolutionary divergence in codon-usage constraints among major plant lineages. Further, the codon interaction network shows global connectivity among codons and reflects the structural relationships underlying codon usage preferences (Figure 8B). The highly connected network is represented by thick lines, indicating codons with strong co-usage tendencies across the species and lineages. The presence of mixed edges with strong (blue) and weak (red) networks showed a complex interplay of evolutionary forces where some codons evolved in a highly interdependent manner while others evolved independently. The network shows that codon usage depends not only on codon preference but on co-evolutionary relationships in the translational system.

**Figure 9.**
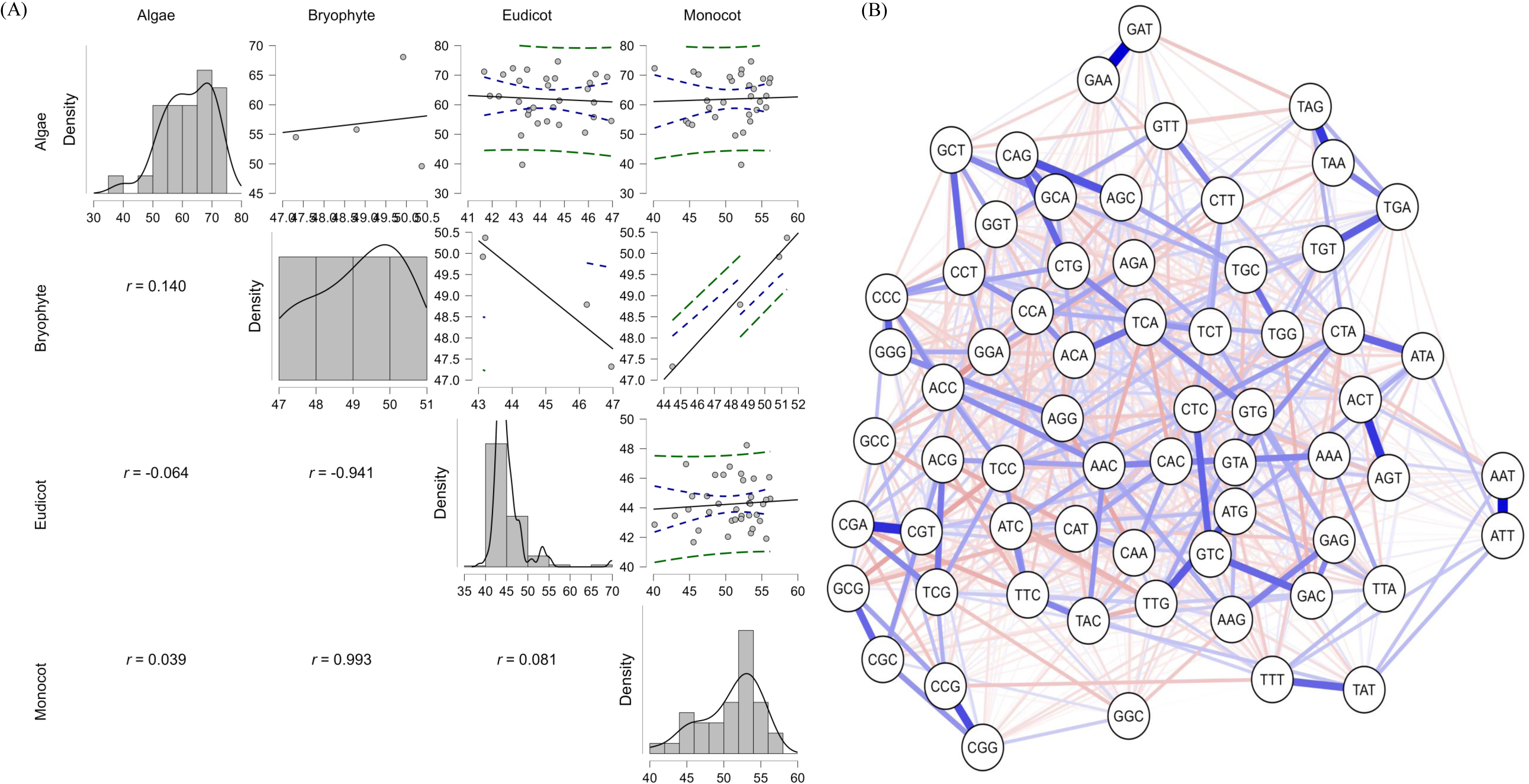
Comparative codon-usage correlations and codon-interaction network architecture across major plant lineages. Panel (A) represents pairwise correlation matrices for codon usage bias in the lower (algae, bryophytes) and the higher (monocots and eudicots) plants. The upper-right panel shows scatterplots with a fitted regression line and 95% confidence intervals. The lower part of the left panel represents Pearson’s correlation coefficient *r*. The diagonal panels represent the lineage-specific variation in codon-usage patterns. Panel (B) shows the network plot of codons generated from the full dataset of codons. Thick blue edges represents strong and positive correlations, whereas red edges show weaker and negative associations. The network configuration emphasizes structural grouping among codons possessing analogous functional or compositional characteristics.

## Discussion

The present genome-wide codon usage bias of 262 plant genomes reveals that the codon usage bias in the plant kingdom is neither stochastic nor uniform. Instead, it reflects a deeply conserved, lineage-specific, and compositionally constrained evolutionary architecture. By integrating codon usage frequencies, synonymous codon usage patterns, codon-anticodon pairing, codon composition, and machine learning analysis, the study uncovered a multi-layered regulatory logic that uncovers mutational dynamics, phylogenetic divergence across the plant lineage, and translational efficiency. Mutational pressure and nucleotide composition were the primary determinants, strongly shaped by GC3 constraints. The positive correlation between GC1,2 and GC3 and the inverse relationship between ENC and GC3 proved that the synonymous sites evolved under a dominant mutational regime. The correlation slope of approximately 0.61 in the neutrality plot (Figure 6D) showed that while mutation was the primary force, selection modulated the base composition at nonsynonymous positions. This dual influence aligns with codon-level evolutionary patterns, where mutational biases shape the compositional landscape and selection pressures. Codon usage was subsequently fine-tuned to increase translational efficiency and fidelity.

It was found that the AT-rich codons were more prominent in higher plants, whereas lower plants, including algae and Bryophytes were dominated by GC-rich codons. This reflected a deep evolutionary divergence in genome architecture shaped by codon usage bias. Wu et al. (2025) reported that genomic architecture evolved through this divergence principle ^36^. The transition from AT-rich codons in lower plants to GC-rich codons in higher plants indicates a shift in metabolic constraints and ecological adaptations ^37,38^. The dominance of preferred codons in highly expressed genes, clustering of synonymous codons in dendrogram and PCA analysis, and elevated CAI values across the species collectively provide strong evidence of the role of translational selection in shaping codon evolution ^14,39^. This was further supported by the codon-anticodon mapping. The clustering of the high-frequency codons with their cognate anticodons along with the presence of structured and near-cognate interactions highlights the co-evolution of codon preference and anticodon availability ^40,41^. This codon-anticodon evolution minimizes the translational errors and optimizes the ribosomal outputs. Further, lineage-specific codon-anticodon pairing, particularly in monocots compared with eudicots, underscores the evolutionary plasticity of protein translation systems.

A lineage-dependent stratification was also evident in ENC distributions, PR2 plots, and RSCU heatmaps. This underscores the role of phylogenetic history in shaping the codon usage in the plant kingdom ^42,43^. The algae and bryophytes exhibited heterogeneous and extreme codon usage bias, consistent with their mutational environment. In contrast, the angiosperm lineage provided more constrained and homogeneous codon usage patterns. This shows a strong selection pressure and stable genome organization ^44,45^. The strong codon usage bias separating higher and lower plants suggests that codon usage bias can serve as a molecular marker of evolutionary transitions. The presence of A/T-ending codons in higher plants and G/C-ending codons in early diverging lineages reflects the broader evolutionary shift from GC-rich to AT-rich genomes during adaptation to terrestrial ecosystems ^37,46^.

In the machine learning analysis, we found CTC codon as the primary predictor in the decision-tree model along with a few subsets of codons across the accuracy and impurity-based rankings. This shows that codon usage bias is not distributed evenly across the genome; instead, a few small numbers of codons carry the disproportionate evolutionary and translational information ^47,48^. The neural network further explained that these codons act as central hubs within a non-linear and interconnected codon landscape. This further proves that codon usage is governed by hierarchical constraints where a few codons exert strong influence on the translational dynamics while others remain functionally redundant ^41,42,47^.

The energy constraints and molecular economy study showed a narrow distribution across the species despite vast differences in the genome size, codon composition, and evolutionary history. This suggests that translational energy optimization is a conserved constraint that requires ∼ 30 ATP towards the synthesis of a codon ^49^. This shows plants have maintained a tightly regulated and conserved energy budget for protein synthesis. This reflects the high metabolic cost of protein translation and the requirement for efficient resource allocations ^30,50^. However, minor lineage-specific variations are driven by mutational and selective pressures. Selective forces strongly stabilize the energetic cost of protein translation by helping preserve metabolic homeostasis. Further, the analysis of codon-usage correlation matrices’ network structure reflects that the codons do not evolve independently ^51^. Instead, they form interconnected modules strongly influenced by the shared mutational bias, tRNA availability, and translational constraints. The coexistence of strong and weak network connections hints at a complex interplay of co-evolutionary dynamics, where certain codons are tightly coordinated with a few others follow independent evolutionary trajectories.

## Conclusions

This study revealed that codon usage bias is an emergent property of the protein translation system shaped by synonymous codon families and genome-wide composition constraints. The study showed that codon usage bias in the plant kingdom is a multifactorial phenomenon shaped by the interplay between mutational pressure and translational selection. The study also revealed that codon usage is not merely a reflection of genome composition but a dynamic evolutionarily optimized system that regulates accurate protein synthesis. The insight gained from this study can have broad implications for plant molecular evolution and the optimization of transgene expression across different plant systems.

## Declaration

**Clinical Trial Number** Not applicable **Acknowledgement** Not available

## Conflict of interest

There is no conflict of interest to declare.

## Compliance with Ethics Requirements

Ethical approval is not applicable in this study. The study used nucleotide sequences downloaded from the publicly available NCBI database. No plant, animal, or microbial samples directly or indirectly used in this study. Python code was used from code libraries and Codex.

## Data availability

Not applicable

## Ethics approval and consent to participate

Not applicable

## Consent for publication

Not applicable

## Author contribution

TKM: conceptualization, methodology, software, validation, formal analysis, investigation, resources, data curation, writing original draft, writing and editing, visualization, supervision, and project administration; SKP: revised and edited the manuscript; HT: analyzed the data and revised the manuscript; PK: analyzed data; JK: Interpreted the data and revised the manuscript.

## Funding

Not available

## Supplementary materials

**Supplementary Figure 1.** Synonymous codon usage frequencies across the plant kingdom. Bars represent mean codon frequencies (%) across all analyzed plant species, with error bars indicating variability among species. Codon usage is shown separately for each amino acid family.

**Supplementary Figure 2.**

Purine-pyrimidine wise nucleotide position in the codon usage of the plant kingdom.

**Supplementary File 1.**

Codon-adaptation index of the codons of the plant kingdom used in this study.

**Supplementary Table 1.**

Species details and their accession numbers used during this study. The table also contains the number of CDS sequences and the number of codons used in this study.

**Supplementary Table 2.**

The highest and lowest codon-usage percentage of individual species.

## Notes

### Competing Interest Statement

The authors have declared no competing interest.

