## Supplementary figures and images for "Evolutionary Stratification of Codon Usage Bias In Plants Arises from GC3 Composition and Translational Optimization"

### Supplementary Figure 1

#### Slide 1
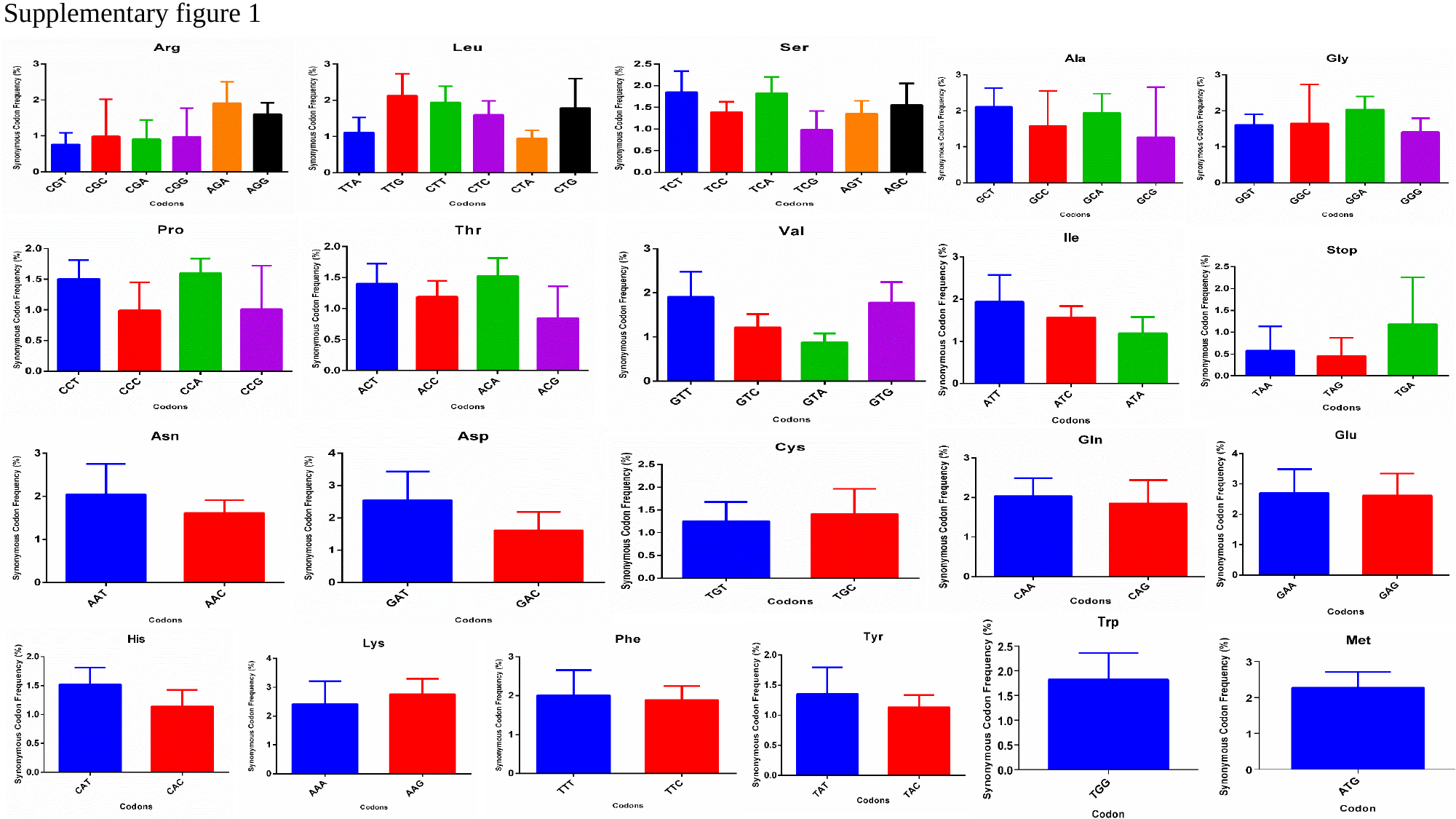

Supplementary figure 1

### Supplementary Figure 2

#### Slide 1
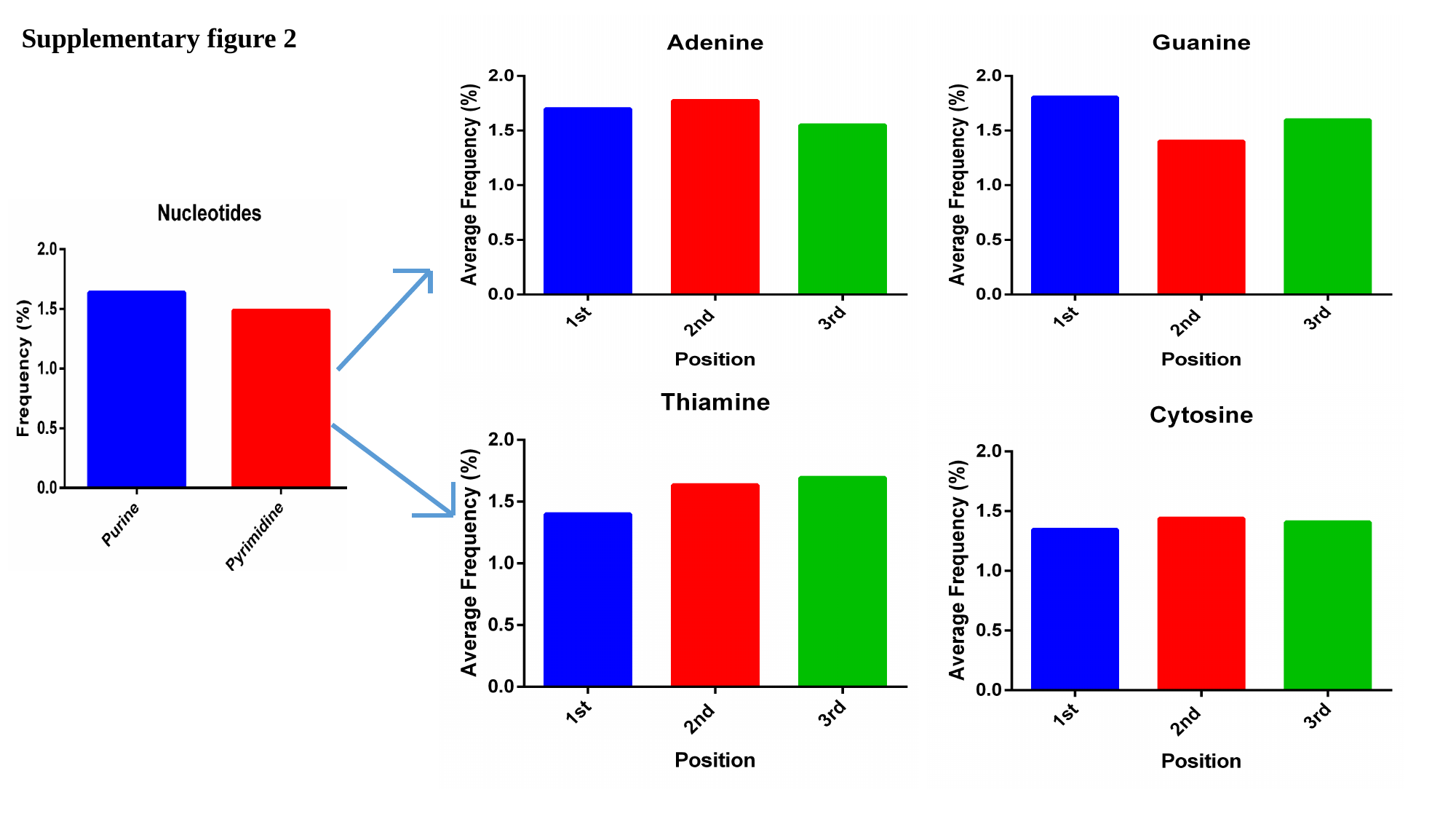

Supplementary figure 2
