## Supplementary Table 1 for "Evolutionary Stratification of Codon Usage Bias In Plants Arises from GC3 Composition and Translational Optimization"

Supplementary Table 1. Species details and their accession numbers used during this study. The table also contain number of CDS sequences and number of codons used in this study.

| **Species** | **Assembly/Accession Number** | **No. of CDS Sequences** | **Total No. of Codons** |
| --- | --- | --- | --- |
| *Abrus precatorius* | GCF_003935025.1 | 40048 | 19920610 |
| *Acer yangbiense* | GCA_008009225.1 | 28308 | 12338579 |
| *Actinidia chinensis* | GCA_003024255.1 | 33117 | 14111084 |
| *Actinidia rufa* | GCA_014362265.1 | 52349 | 19065849 |
| *Aegilops tauschii* | GCA_001957025.1 | 258680 | 88004792 |
| *Amborella trichopoda* | GCA_000471905.1 | 26846 | 8467095 |
| *Anacardium occidentale* | Phytozome | 82170 | 34149143 |
| *Ananascomosus* | GCA_001661175.1 | 35775 | 16805721 |
| *Apium graveolens* | GCA_009905375.1 | 1660 | 566725 |
| *Apostasia shenzhenica* | GCA_002786265.1 | 21743 | 7967114 |
| *Aquilegia coerulea* | GCA_002738505.1 | 41603 | 17920131 |
| *Arabidopsis halleri* | Phytozome | 34087 | 13768169 |
| *Arabidopsis lyrata* | GCA_000004255.1 | 39161 | 11804251 |
| *Arabidopsisthaliana* | GCA_000001735.2 | 48265 | 20881456 |
| *Arabis alpina* | GCA_000733195.1 | 23286 | 9131774 |
| *Arabis nemorensis* | GCA_902206195.3 | 28841 | 10676198 |
| *Arachis duranensis* | GCA_000817695.2 | 52826 | 22819450 |
| *Arachis hypogaea* | GCA_003086295.2 | 100775 | 46083242 |
| *Arachis ipaensis* | GCA_000816755.2 | 57621 | 24226039 |
| *Asparagus officinalis* | GCA_001876935.1 | 36763 | 15535738 |
| *Auxenochlorella protothecoide* | GCA_000733215.1 | 7014 | 2764633 |
| *Bathycoccus prasinos* | GCA_002220235.1 | 7900 | 4123485 |
| *Benincasa hispida* | GCA_009727055.1 | 33043 | 15486726 |
| *Beta vulgaris* | GCA_000511025.2 | 32874 | 10338057 |
| *Boechera stricta* | Phytozome | 29812 | 11925325 |
| *Botryococcus braunii* | Phytozome | 23685 | 8370626 |
| *Brachypodium distachyon* | GCA_000005505.4 | 37892 | 22249324 |
| *Brachypodium x hybridum* | Phytozome | 40182 | 14974746 |
| *Brassica carinata* | GCA_016771965.1 | 97148 | 32541732 |
| *Brassica cretica* | GCA_003260655.2 | 63131 | 18215199 |
| *Brassica napus* | GCA_000751015.1 | 101040 | 33712609 |
| *Brassica oleracea* | GCA_000695525.1 | 56687 | 20569585 |
| *Brassica rapa* | Phytozome | 41025 | 16035873 |
| *Cajanus cajan* | GCA_000340665.1 | 41387 | 18977418 |
| *Camelina sativa* | GCA_000633955.1 | 107481 | 45920204 |
| *Camellia sinensis* | GCA_013676235.1 | 32356 | 12182310 |
| *Cannabis sativa* | GCA_900626175.2 | 33674 | 15403654 |
| *Capsella rubella* | GCA_000375325.1 | 34126 | 15426557 |
| *Capsicum annuum* | GCA_000512255.2 | 35845 | 13032315 |
| *Capsicum baccatum* | GCA_002271885.2 | 35853 | 13450553 |
| *Capsicum chinense* | GCA_002271895.2 | 34974 | 13006454 |
| *Carex littledale* | GCA_011114355.1 | 22441 | 8685207 |
| *Carica papaya* | GCA_000150535.1 | 26103 | 10815624 |
| *Carpinus fangiana* | GCA_006937295.1 | 29499 | 12909669 |
| *Carya illinoinensis* | GCA_016808215.1 | 68161 | 35693735 |
| *Castanea mollissima* | GCA_000763605.2 | 33467 | 13521297 |
| *Cephalotus follicularis* | GCA_001972305.1 | 36667 | 11659493 |
| *Ceratodon purpureus* | GCA_014871385.1 | 40806 | 16203113 |
| *Chara braunii* | GCA_003427395.1 | 35665 | 23488979 |
| *Chenopodium quinoa* | GCA_001683475.1 | 63173 | 28189986 |
| *Chlamydomonas eustigma* | GCA_002335675.1 | 14161 | 8178670 |
| *Chlamydomonas incerta* | GCA_016834605.1 | 16957 | 13646559 |
| *Chlamydomonas reinhardtii* | GCA_000002595.2 | 14488 | 13362550 |
| *Chlamydomonas schloesseri* | GCA_016834595.1 | 16268 | 13160388 |
| *Chlorella sorokiniana* | GCA_002245835.2 | 10384 | 6815625 |
| *Chlorella variabilis* | GCA_000147415.1 | 9780 | 4456440 |
| *Chondrus crispus* | PRJEA78309 | 9603 | 3424143 |
| *Chromochloris zofingiensis* | Phytozome | 15369 | 7400426 |
| *Cicer arietinum* | GCA_000331145.1 | 35754 | 16569770 |
| *Cinnamomum micranthum* | GCA_003546025.1 | 26531 | 12185343 |
| *Citrus clementina* | GCA_000493195.1 | 32586 | 14021931 |
| *Citrus sinensis* | GCA_000695605.1 | 51718 | 18652779 |
| *Citrus unshiu* | GCA_002897195.1 | 37970 | 15899553 |
| *Coccomyxa subellipsoidea* | GCA_000258705.1 | 9839 | 4187896 |
| *Coffea arabica* | GCA_003713225.1 | 67222 | 30079196 |
| *Coffea canephora* | GCA_900059795.1 | 25574 | 10276925 |
| *Coffea eugenioides* | GCA_003713205.1 | 38150 | 18082436 |
| *Coptis chinensis* | GCA_015680905.1 | 40011 | 12856992 |
| *Corchorus capsularis* | GCA_001974805.1 | 29356 | 10123762 |
| *Corchorus olitorius* | GCA_001974825.1 | 35704 | 11159127 |
| *Corymbia citriodora* | GCA_014858505.1 | 45647 | 19782427 |
| *Cucumis melo* | GCA_005549215.1 | 38173 | 14259790 |
| *Cucumis sativus* | GCA_000004075.3 | 30919 | 8765187 |
| *Cucurbita maxima* | GCA_002738345.1 | 42777 | 20412789 |
| *Cucurbita moschata* | GCA_002738365.1 | 43715 | 20980835 |
| *Cucurbita pepo* | GCA_002806865.2 | 25263 | 20372649 |
| *Cuscuta australis* | GCA_003260385.1 | 18157 | 7490641 |
| *Cuscuta campestris* | GCA_900332095.2 | 45474 | 18652802 |
| *Cyanidioschyzon merolae* | ASM623231v1 | 252 | 2510657 |
| *Cynara cardunculus* | GCA_001531365.1 | 38406 | 18981503 |
| *Daucus carota* | GCA_001625215.1 | 44655 | 12712788 |
| *Dendrobium catenatum* | GCA_001605985.2 | 34389 | 15864674 |
| *Dichanthelium oligosanthes* | GCA_001633215.2 | 26368 | 9740302 |
| *Digitaria exilis* | GCA_015342445.1 | 61983 | 27775825 |
| *Dioscorea cayenensis* | GCA_009730915.2 | 33298 | 14897198 |
| *Dioscorea rotundata* | GCF_009730915.1 | 19023 | 8109698 |
| *Dorcoceras hygrometricum* | GCA_001598015.1 | 47778 | 15189804 |
| *Dunaliella salina* | GCA_002284615.2 | 18740 | 7872404 |
| *Durio zibethinus* | GCA_002303985.1 | 63007 | 31840756 |
| *Edaphochlamys debaryana* | GCA_016858145.1 | 20450 | 15246466 |
| *Elaeis guineensis* | GCA_000442705.1 | 43551 | 20623029 |
| *Ensete ventricosum* | GCA_000818735.3 | 58438 | 13171749 |
| *Eragrostis curvula* | GCA_007726485.1 | 55182 | 13171749 |
| *Eragrostis tef* | ASM97063v1 | 41555 | 16268882 |
| *Erythranthe guttata* | GCA_000504015.1 | 31861 | 14340456 |
| *Eucalyptus grandis* | GCA_000612305.1 | 46920 | 24684301 |
| *Eutrema salsugineum* | GCA_000478725.1 | 33637 | 15105782 |
| *Fragaria vesca* | Phytozome | 31387 | 14541202 |
| *Galdieria sulphuraria* | ASM34128v1 | 7174 | 3012469 |
| *Genlisea aurea* | GCA_000441915.1 | 17685 | 5697418 |
| *Glycine max* | Ensemble | 76522 | 37588895 |
| *Glycine soja* | GCA_000722935.2 | 50399 | 18475984 |
| *Gonium pectorale* | GCA_001584585.1 | 16290 | 8150529 |
| *Gossypium arboreum* | GCA_000787975.1 | 33609 | 9425236 |
| *Gossypium aridum* | GCA_013487665.1 | 26546 | 8514527 |
| *Gossypium armourianum* | GCA_013677265.1 | 23673 | 8494193 |
| *Gossypium australe* | GCA_005393395.2 | 38281 | 14731462 |
| *Gossypium barbadense* | GCA_008761655.1 | 108363 | 46191606 |
| *Gossypium darwinii* | GCA_007990325.1 | 97407 | 38830197 |
| *Gossypium davidsonii* | GCA_013677245.1 | 29468 | 10991130 |
| *Gossypium gossypioides* | GCA_013467495.1 | 21018 | 7999915 |
| *Gossypium harknessii* | GCA_013677255.1 | 27624 | 9012849 |
| *Gossypium hirsutum* | GCA_000987745.1 | 90927 | 40108774 |
| *Gossypium klotzschianum* | GCA_013677235.1 | 30394 | 11059532 |
| *Gossypium laxum* | GCA_013511315.1 | 25577 | 9688534 |
| *Gossypium lobatum* | GCA_013467485.1 | 27087 | 8961513 |
| *Gossypium mustelinum* | GCA_007990455.1 | 106487 | 43711902 |
| *Gossypium raimondii* | GCA_000327365.1 | 59057 | 32979788 |
| *Gossypium schwendimanii* | GCA_013677275.1 | 29079 | 11374820 |
| *Gossypium tomentosum* | GCA_007990485.1 | 112713 | 46891773 |
| *Gossypium trilobum* | GCA_013467465.1 | 27591 | 10636279 |
| *Haematococcus lacustris* | GCA_011766145.1 | 28279 | 5222227 |
| *Handroanthus impetiginosus* | GCA_002762385.1 | 30271 | 11293274 |
| *Helianthus annuus* | GCA_002127325.2 | 75695 | 18113290 |
| *Helicosporidium sp.* | GCA_000690575.1 | 6033 | 1651430 |
| *Herrania umbratica* | GCA_002168275.2 | 27748 | 13475720 |
| *Hevea brasiliensis* | GCA_010458925.1 | 44146 | 16775789 |
| *Hibiscus syriacus* | GCA_006381635.2 | 81925 | 33103637 |
| *Hordeum vulgare* | GCA_902500625.1 | 236194 | 85244531 |
| *Ipomoea nil* | GCA_001879475.1 | 51054 | 44173 |
| *Ipomoea triloba* | ASM357664v1 | 47019 | 20742621 |
| *Jatropha curcas* | GCA_014843425.1 | 29586 | 14084402 |
| *Juglans regia* | GCA_001411555.2 | 45959 | 22099182 |
| *Kalanchoe fedtschenkoi* | Phytozome | 45190 | 18509074 |
| *Kalanchoe laxiflora* | Phytozome | 69177 | 30253473 |
| *Kingdonia uniflora* | GCA_014058105.1 | 43300 | 13809631 |
| *Klebsormidium nitens* | GCA_000708835.1 | 16283 | 8787539 |
| *Lactuca saligna* | GCA_902860255.1 | 45452 | 16922724 |
| *Lactuca sativa* | GCA_002870075.1 | 45242 | 19759541 |
| *Leersia perrieri* | Ensemble | 38960 | 17378001 |
| *Linum usitatissimum* | Phytozome | 43484 | 17374241 |
| *Lupinus albus* | GCA_010261695.1 | 47603 | 18939592 |
| *Lupinus angustifolius* | GCA_001865875.1 | 52821 | 14210439 |
| *Macleaya cordata* | GCA_002174775.1 | 21911 | 9192500 |
| *Malus baccata* | GCA_006547085.1 | 45900 | 16274185 |
| *Malus domestica* | Phytozome | 40624 | 15853286 |
| *Manihot esculenta* | GCA_001659605.1 | 43286 | 16979662 |
| *Marchantia polymorpha* | Phytozome | 24674 | 10963140 |
| *Medicago truncatula* | Phytozome | 29448 | 19930842 |
| *Micractinium conductrix* | GCA_002245815.2 | 10070 | 6540083 |
| *Micromonas commoda* | GCA_000090985.2 | 10137 | 4906972 |
| *Micromonas pusilla* | GCA_000151265.1 | 10242 | 4605733 |
| *Microthlaspi erraticum* | GCA_902728155.2 | 49768 | 17774892 |
| *Mikania micrantha* | GCA_009363875.1 | 46329 | 19197172 |
| *Mimulus guttatus* | Phytozome | 33573 | 13654273 |
| *Miscanthus lutarioriparius* | GCA_904845875.1 | 68287 | 27677044 |
| *Miscanthus sinensis* | Phytozome | 89486 | 33209433 |
| *Momordica charantia* | GCA_001995035.1 | 28666 | 13384679 |
| *Monoraphidium neglectum* | GCA_000611645.1 | 16755 | 5836502 |
| *Morella rubra* | GCA_003952965.2 | 29351 | 11191982 |
| *Morus notabilis* | GCA_000414095.2 | 27648 | 16813533 |
| *Musa acuminata* | GCA_000313855.2 | 47707 | 12585400 |
| *Musa balbisiana* | GCA_004837865.1 | 33021 | 13407392 |
| *Nelumbo nucifera* | GCA_000365185.2 | 38191 | 18623218 |
| *Nicotiana attenuata* | Ensemble | 28935 | 12378926 |
| *Nicotiana sylvestris* | GCA_000393655.1 | 48160 | 20526275 |
| *Nicotiana tabacum* | GCA_000715135.1 | 84255 | 34958126 |
| *Nicotiana tomentosiformis* | GCA_000390325.2 | 50113 | 21963849 |
| *Nymphaea colorata* | GCA_008831285.1 | 33500 | 16412348 |
| *Nymphaea thermarum* | GCA_011799765.1 | 25461 | 10353006 |
| *Nyssa sinensis* | GCA_008638375.1 | 36241 | 14291804 |
| *Olea europaea* | Ensemble | 50678 | 17584084 |
| *Oryza barthii* | Ensemble | 41595 | 16853621 |
| *Oryza brachyantha* | Ensemble | 32037 | 11135617 |
| *Oryza glaberrima* | Ensemble | 33164 | 12097072 |
| *Oryza glumipatula* | Ensemble | 46893 | 19359968 |
| *Oryza sativa indica* | ASM465v1 | 40745 | 15146969 |
| *Oryza longistaminata* | Ensemble | 31686 | 11577073 |
| *Oryza meridionalis* | Ensemble | 43455 | 18403462 |
| *Oryza nivara* | Ensemble | 48360 | 20335202 |
| *Oryza punctata* | Ensemble | 41060 | 17827430 |
| *Oryza rufipogon* | Ensemble | 47411 | 19633269 |
| *Oryza sativa* | IRGSP-1.0 | 42246 | 14018071 |
| *Ostreococcus lucimarinus* | ASM9206v1 | 7603 | 3075822 |
| *Ostreococcus tauri* | GCA_000214015.2 | 7766 | 3573773 |
| *Panicum hallii* | Ensemble | 44192 | 18118182 |
| *Panicum miliaceum* | GCA_003046395.2 | 55964 | 21862123 |
| *Panicum virgatum* | GCA_016808335.1 | 129942 | 58265831 |
| *Papaver somniferum* | GCA_003573695.1 | 84179 | 35992719 |
| *Parasponia andersonii* | GCA_002914805.1 | 37227 | 11059905 |
| *Phalaenopsis equestris* | GCA_001263595.1 | 29894 | 13085782 |
| *Phaseolus vulgaris* | PhaVulg1 | 32720 | 13732964 |
| *Phoenix dactylifera* | GCA_009389715.1 | 48801 | 23300984 |
| *Physcomitrella patens* | Ensemble | 86669 | 38906629 |
| *Pistacia vera* | Ensemble | 31784 | 12704908 |
| *Populus alba* | GCA_005239225.1 | 46966 | 23277683 |
| *Populus deltoides* | GCA_015852605.1 | 44265 | 18589863 |
| *Populus euphratica* | GCA_000495115.1 | 49760 | 24591814 |
| *Populus trichocarpa* | Ensemble | 73102 | 30267150 |
| *Prosopis alba* | GCA_004799145.1 | 57572 | 25824631 |
| *Prunus armeniaca* | GCA_903112645.1 | 30573 | 10602780 |
| *Prunus avium* | GCA_002207925.1 | 35009 | 12920160 |
| *Prunus dulcis* | GCA_902201215.1 | 33326 | 16453804 |
| *Prunus mume* | GCA_000346735.1 | 29705 | 13783885 |
| *Prunus persica* | GCA_000346465.2 | 32595 | 21638326 |
| *Prunus yedoensis* | GCA_002966975.2 | 41294 | 13134476 |
| *Punica granatum* | GCA_007655135.2 | 36608 | 17742114 |
| *Pyrus x bretschneideri* | GCA_000315295.1 | 47086 | 22153368 |
| *Quercus lobata* | GCA_001633185.2 | 53228 | 26033433 |
| *Quercus suber* | GCA_002906115.1 | 59614 | 27643487 |
| *Raphanus sativus* | GCA_000801105.2 | 61216 | 25228923 |
| *Raphidocelis subcapitata* | GCA_003203535.1 | 13383 | 7510444 |
| *Rhamnella rubrinervis* | GCA_007844105.2 | 27165 | 11450813 |
| *Rhododendron simsii* | GCA_014282245.1 | 32999 | 14175587 |
| *Rhododendron williamsianum* | GCA_009746105.1 | 23548 | 9672699 |
| *Ricinus communis* | GCA_000151685.2 | 28584 | 13314890 |
| *Rosa chinensis* | GCA_002994745.1 | 45097 | 21728915 |
| *Saccharum spontaneum* | GCA_003544955.1 | 53284 | 20732408 |
| *Salix brachista* | GCA_009078335.1 | 30209 | 13198070 |
| *Salix dunnii* | GCA_015731905.1 | 29822 | 12885851 |
| *Salvia splendens* | GCA_004379255.1 | 53354 | 22961551 |
| *Scenedesmus sp.* | GCA_014080715.1 | 17398 | 7082477 |
| *Selaginella moellendorffii* | GCA_000143415.2 | 34825 | 13726751 |
| *Senna tora* | GCA_014851425.1 | 45268 | 14353185 |
| *Sesamum indicum* | GCA_000512975.1 | 35410 | 16628902 |
| *Setaria italica* | GCA_000263155.2 | 35844 | 16918526 |
| *Setaria viridis* | GCA_005286985.1 | 39206 | 18608154 |
| *Sinapis alba* | GCA_012274485.2 | 52686 | 20255730 |
| *Solanum lycopersicum* | Phytozome | 34429 | 12189341 |
| *Solanum pennellii* | GCA_001406875.2 | 38023 | 17963715 |
| *Solanum tuberosum* | GCA_000226075.1 | 37966 | 16949304 |
| *Sorghum bicolor* | GCA_000003195.3 | 47110 | 19286451 |
| *Spatholobus suberectus* | GCA_004329165.1 | 31106 | 11450383 |
| *Sphagnum fallax* | Phytozome | 26939 | 12143752 |
| *Spinacia oleracea* | GCA_002007265.1 | 32794 | 14915256 |
| *Spirodela polyrhiza* | Phytozome | 19623 | 7247698 |
| *Striga asiatica* | GCA_008636005.1 | 33426 | 12366431 |
| *Syzygium oleosum* | GCA_900635055.1 | 38158 | 17678522 |
| *Tarenaya hassleriana* | GCA_000463585.1 | 41094 | 18822592 |
| *Tetrabaena socialis* | GCA_002891735.1 | 14296 | 5805409 |
| *Tetracentron sinense* | GCA_015143295.1 | 32739 | 14232443 |
| *Thalictrum thalictroides* | GCA_013358455.1 | 33624 | 10525461 |
| *Theobroma cacao* | GCA_000403535.1 | 47110 | 19420974 |
| *Trebouxia sp.* | GCA_008636185.1 | 13918 | 5659886 |
| *Trema orientale* | GCA_002914845.1 | 35849 | 11293853 |
| *Trifolium pratense* | Ensemble | 41270 | 14114820 |
| *Trifolium subterraneum* | GCA_001742945.1 | 42059 | 15474885 |
| *Tripterygium wilfordii* | GCA_013401445.1 | 48101 | 22731305 |
| *Triticum aestivum* | IWGSC RefSeq v1.1 | 133346 | 59205793 |
| *Triticum dicoccoides* | GCA_002162155.1 | 295286 | 38245508 |
| *Triticum turgidum* | Ensemble | 196105 | 90475063 |
| *Triticum urartu* | ASM34745v1 | 33482 | 12164580 |
| *Vanilla planifolia* | GCA_016413885.1 | 29180 | 9584827 |
| *Vigna angularis* | Vigan1.1 | 33860 | 12601663 |
| *Vigna radiata* | Ensemble | 23181 | 9913023 |
| *Vigna unguiculata* | GCA_004118075.1 | 41173 | 19723187 |
| *Vitis riparia* | GCA_004353265.1 | 39906 | 19821049 |
| *Vitis vinifera* | Phytozome | 29927 | 10783958 |
| *Volvox carteri* | GCA_000143455.1 | 14436 | 8215905 |
| *Zea mays* | PRJNA10769 | 131318 | 63461672 |
| *Ziziphus jujuba* | GCA_000826755.1 | 43574 | 20144373 |
| *Zostera marina* | GCA_001185155.1 | 20648 | 8030995 |
