## Supplementary Table 2 for "Evolutionary Stratification of Codon Usage Bias In Plants Arises from GC3 Composition and Translational Optimization"

Supplementary Table 2. The highest and lowest percentage of codon usage of individual species.

| **Codons** | **Highest %** | **Species with Highest %** | **Lowest %** | **Species With Lowest %** |
| --- | --- | --- | --- | --- |
| AAA | 3.930 | *Bathycoccus prasinos* | 0.169 | *Edaphochlamys debaryana* |
| AAC | 2.989 | *Cicer arietinum* | 0.358 | *Chlorella sorokiniana* |
| AAG | 3.546 | *Spirodela polyrhiza* | 0.369 | *Chlorella sorokiniana* |
| AAT | 3.187 | *Ipomoea nil* | 0.144 | *Raphidocelis subcapitata* |
| ACA | 2.620 | *Cicer arietinum* | 0.393 | *Gonium pectorale* |
| ACC | 2.595 | *Cicer arietinum* | 0.655 | *Chlorella sorokiniana* |
| ACG | 3.758 | *Micromonas pusilla* | 0.367 | *Aquilegia coerulea* |
| ACT | 2.516 | *Cicer arietinum* | 0.317 | *Tetrabaena socialis* |
| AGA | 3.042 | *Cucurbita maxima* | 0.095 | *Gonium pectorale* |
| AGC | 4.561 | *Scenedesmus sp* | 0.916 | *Aquilegia coerulea* |
| AGG | 3.079 | *Chara braunii* | 0.562 | *Chlamydomonas reinhardtii* |
| AGT | 2.286 | *Cicer arietinum* | 0.248 | *Tetrabaena socialis* |
| ATA | 2.284 | *Cicer arietinum* | 0.101 | *Chlorella sorokiniana* |
| ATC | 2.407 | *Genlisea aurea* | 0.331 | *Chlorella sorokiniana* |
| ATG | 3.478 | *Rosa chinensis* | 0.353 | *Chlorella sorokiniana* |
| ATT | 3.101 | *Ipomoea nil* | 0.170 | *Raphidocelis subcapitata* |
| CAA | 3.255 | *Chlorella sorokiniana* | 0.420 | *Gonium pectorale* |
| CAC | 2.649 | *Chlorella sorokiniana* | 0.712 | *Aquilegia coerulea* |
| CAG | 5.338 | *Scenedesmus sp* | 0.756 | *Ipomoea nil* |
| CAT | 2.513 | *Chlorella sorokiniana* | 0.284 | *Gonium pectorale* |
| CCA | 2.610 | *Chlorella variabilis* | 0.603 | *Gonium pectorale* |
| CCC | 3.427 | *Edaphochlamys debaryana* | 0.379 | *Bathycoccus prasinos* |
| CCG | 4.683 | *Raphidocelis subcapitata* | 0.323 | *Aquilegia coerulea* |
| CCT | 2.993 | *Chlorella sorokiniana* | 0.545 | *Ostreococcus lucimarinus* |
| CGA | 4.186 | *Ostreococcus lucimarinus* | 0.340 | *Gonium pectorale* |
| CGC | 6.085 | *Raphidocelis subcapitata* | 0.312 | *Thalictrum thalictroides* |
| CGG | 5.365 | *Chlamydomonas incerta* | 0.368 | *Glycine soja* |
| CGT | 2.792 | *Micromonas pusilla* | 0.374 | *Durio zibethinus* |
| CTA | 1.624 | *Quercus suber* | 0.228 | *Raphidocelis subcapitata* |
| CTC | 3.015 | *Spirodela polyrhiza* | 0.844 | *Chromochloris zofingiensis* |
| CTG | 6.257 | *Gonium pectorale* | 0.522 | *Ipomoea nil* |
| CTT | 2.647 | *Aquilegia coerulea* | 0.476 | *Raphidocelis subcapitata* |
| GAA | 4.146 | *Bathycoccus prasinos* | 0 | *Coffea arabica* |
| GAC | 3.986 | *Gonium pectorale* | 0.746 | *Durio zibethinus* |
| GAG | 5.065 | *Gonium pectorale* | 1.011 | *Ipomoea nil* |
| GAT | 4.064 | *Lactuca saligna* | 0 | *Coffea arabica* |
| GCA | 5.445 | *Scenedesmus sp* | 0.953 | *Ipomoea nil* |
| GCC | 6.039 | *Chlamydomonas reinhardtii* | 0.556 | *Galdieria sulphuraria* |
| GCG | 8.424 | *Raphidocelis subcapitata* | 0.369 | *Durio zibethinus* |
| GCT | 5.212 | *Chlorella sorokiniana* | 0.930 | *Ipomoea nil* |
| GGA | 3.913 | *Chlorella sorokiniana* | 0.671 | *Tetrabaena socialis* |
| GGC | 6.665 | *Chlamydomonas schloesseri* | 0 | *Chlamydomonas reinhardtii* |
| GGG | 3.250 | *Edaphochlamys debaryana* | 0.592 | *Galdieria sulphuraria* |
| GGT | 2.439 | *Chromochloris zofingiensis* | 0.786 | *Raphidocelis subcapitata* |
| GTA | 1.347 | *Kingdonia uniflora* | 0.144 | *Raphidocelis subcapitata* |
| GTC | 2.355 | *Micromonas pusilla* | 0.661 | *Chlorella variabilis* |
| GTG | 4.432 | *Chlamydomonas reinhardtii* | 0.516 | *Ipomoea nil* |
| GTT | 2.911 | *Aquilegia coerulea* | 0.363 | *Auxenochlorella protothecoide* |
| TAA | 2.080 | *Ipomoea nil* | 0.031 | *Tetrabaena socialis* |
| TAC | 1.765 | *Gonium pectorale* | 0.258 | *Chlorella sorokiniana* |
| TAG | 1.371 | *Ipomoea nil* | 0.027 | *Genlisea aurea* |
| TAT | 3.085 | *Ipomoea nil* | 0.096 | *Raphidocelis subcapitata* |
| TCA | 2.453 | *Ipomoea nil* | 0.367 | *Gonium pectorale* |
| TCC | 2.181 | *Spirodela polyrhiza* | 0.486 | *Chlorella sorokiniana* |
| TCG | 3.507 | *Micromonas pusilla* | 0.325 | *Chromochloris zofingiensis* |
| TCT | 2.621 | *Aquilegia coerulea* | 0.345 | *Tetrabaena socialis* |
| TGA | 4.308 | *Cucurbita maxima* | 0.065 | *Chlamydomonas reinhardtii* |
| TGC | 3.860 | *Scenedesmus sp* | 0.607 | *Cicer arietinum* |
| TGG | 3.706 | *Chlorella variabilis* | 0.554 | *Cicer arietinum* |
| TGT | 2.542 | *Cucurbita maxima* | 0.144 | *Gonium pectorale* |
| TTA | 1.783 | *Trifolium pratense* | 0.054 | *Gonium pectorale* |
| TTC | 2.739 | *Genlisea aurea* | 0.330 | *Chlorella sorokiniana* |
| TTG | 3.591 | *Durio zibethinus* | 0.277 | *Chlorella sorokiniana* |
| TTT | 4.432 | *Ipomoea nil* | 0.197 | *Chlamydomonas incerta* |
